# Defective interferon signaling in the circulating monocytes of type 2 diabetic mice

**DOI:** 10.1101/2024.06.03.597050

**Authors:** Shunsuke Omodaka, Yuya Kato, Yoshimichi Sato, Jaime Falcone-Juengert, Hongxia Zhang, Atsushi Kanoke, Walter L. Eckalbar, Hidenori Endo, Christine L. Hsieh, Dvir Aran, Jialing Liu

**Affiliations:** Department of Neurological Surgery, University of California San Francisco, San Francisco, CA 94158; San Francisco VA Health Care System, San Francisco, CA 94158, 94121; Department of Neurosurgery, Tohoku University Graduate School of Medicine, Sendai 9808574, Japan; Department of Medicine, University of California San Francisco, San Francisco, CA 94117; Department of Medicine, Division of Rheumatology, University of California San Francisco, San Francisco, CA, USA; Faculty of Biology, Technion, Israel Institute of Technology, Haifa, Israel; The Taub Faculty of Computer Science, Technion-Israel Institute of Technology, Haifa, Israel

**Keywords:** single-cell RNA sequencing, bulk RNA sequencing, inflammation, coagulation, Interferon, STAT1, TLR4, preconditioning, neuroprotection, immunosuppression, MHC

## Abstract

Type 2 diabetes mellitus (T2DM) is associated with poor outcome after stroke. Peripheral monocytes play a critical role in the secondary injury and recovery of damaged brain tissue after stroke, but the underlying mechanisms are largely unclear. To investigate transcriptome changes and molecular networks across monocyte subsets in response to T2DM and stroke, we performed single-cell RNA-sequencing (scRNAseq) from peripheral blood mononuclear cells and bulk RNA-sequencing from blood monocytes from four groups of adult mice, consisting of T2DM model *db/db* and normoglycemic control *db/+* mice with or without ischemic stroke. Via scRNAseq we found that T2DM expands the monocyte population at the expense of lymphocytes, which was validated by flow cytometry. Among the monocytes, T2DM also disproportionally increased the inflammatory subsets with Ly6C+ and negative MHC class II expression (MO.6C+II−). Conversely, monocytes from control mice without stroke are enriched with steady-state classical monocyte subset of MO.6C+II+ but with the least percentage of MO.6C+II− subtype. Apart from enhancing inflammation and coagulation, enrichment analysis from both scRNAseq and bulk RNAseq revealed that T2DM specifically suppressed type-1 and type-2 interferon signaling pathways crucial for antigen presentation and the induction of ischemia tolerance. Preconditioning by lipopolysaccharide conferred neuroprotection against ischemic brain injury in *db/+* but not in *db/db* mice and coincided with a lesser induction of brain Interferon-regulatory-factor-3 in the brains of the latter mice. Our results suggest that the increased diversity and altered transcriptome in the monocytes of T2DM mice underlie the worse stroke outcome by exacerbating secondary injury and potentiating stroke-induced immunosuppression.

**Significance Statement:** The mechanisms involved in the detrimental diabetic effect on stroke are largely unclear. We show here, for the first time, that peripheral monocytes have disproportionally altered the subsets and changed transcriptome under diabetes and/or stroke conditions. Moreover, genes in the IFN-related signaling pathways are suppressed in the diabetic monocytes, which underscores the immunosuppression and impaired ischemic tolerance under the T2DM condition. Our data raise a possibility that malfunctioned monocytes may systemically and focally affect the host, leading to the poor outcome of diabetes in the setting of stroke. The results yield important clues to molecular mechanisms involved in the detrimental diabetic effect on stroke outcome.

## Introduction

Type 2 diabetes mellitus (T2DM) is not only an established risk factor for ischemic stroke ^1^, but also associated with poor outcomes after stroke ^2,3^. Peripheral monocytes have emerged as a highly plastic and dynamic cellular system that ensures the sensing of and response to brain injury, as well as resolution of inflammatory processes, mirroring the genomic profile of brain injury ^4,5^. Established evidence suggests that monocytes play a critical role in the development of stroke in T2DM through atherogenesis ^6^, yet its role in exacerbating stroke outcome in subjects with T2DM remains unclear.

Multiple classification schemes exist to characterize monocyte subsets depending on the markers and functional implications. Two distinct types of circulating monocytes were most widely documented in mice as classical (Ly6C^hi^) and non-classical (Ly6C^lo^) monocytes ^7^ ^8,9^. The former sense injury and extravasate into tissue where they maturate to macrophages and dendritic cells ^10^, whereas the latter normally patrol the resting vasculature to maintain vessel wall integrity ^11,12^. Monocytes also engage in antigen presentation, during which major histocompatibility complex (MHC) are crucial for the T cells to discriminate between self and nonself antigens. Unlike the universal expression of MHC-I proteins on all nucleated cells, MHC-II molecules are restricted to antigen presenting cells including the monocytes, with either constitutive or inducible expression ^13^ depending on the monocyte subsets ^14^. Ly6C+ monocytes are reported to confer neuroprotection in the brain parenchyma after stroke ^15^ and may mediate host tolerance to ischemia ^16^, yet their role and effectiveness under diabetic condition have not been assessed.

Ischemic tolerance (IT) can be induced by a number of experimental paradigms given prior to, during, or after an ischemic insult, allowing them to “condition” the tissue/organs to become resistant to subsequent ischemic challenge. Exposure to a brief sublethal episode of ischemia or pharmacological agents before a major cerebral ischemia is neuroprotective, a phenomenon called ischemic pre-conditioning (IPC) ^17–20^. IT or IPC is known to decline with age ^21,22^, yet it is unclear whether type 2 diabetes (T2DM)-caused precocious aging also blunts IT and the underlying molecular mechanisms. Preconditioning stimuli to confer neuroprotection to ischemic stroke are best exemplified by brief ischemia itself or Toll-Like Receptor (TLR) agonists, and both notably converge on interferon regulatory factor (IRF)-dependent signaling ^18,19,23^.

Here we used single-cell RNA sequencing (scRNAseq) and bulk RNAseq to investigate gene expression changes in murine peripheral monocytes under conditions of T2DM and stroke, which were validated with additional experimental approaches. The purpose of this study was to determine distinct subsets of monocytes affected by T2DM and/or stroke and specific transcriptome signatures under these conditions, to uncover the molecular mechanisms underlying the detrimental effect of T2DM on stroke-induced secondary injury and dampening effect on preconditioning-induced neuroprotection against stroke injury using the *db/db* mouse model.

## Materials and Methods

### Animals and housing

All animal experiments were approved by the San Francisco VA Medical Center Animal Care and Use Committee. The *db/db* mouse (B6.BKS(D)-Lepr<*db/db*>/J) carrying a mutation in the leptin receptor gene is a well-established rodent model of obesity-induced T2DM ^24^. Heterozygous *db/+* (B6.BKS(D)-Lepr<*db/+*>/J) mice were chosen as the normoglycemic controls. Male *db/db* and *db/+* mice (10–12 weeks old; Jackson Laboratories, ME) were housed 4 per cage on a 12-hour dark/light cycle with access to food and water ad libitum. We performed direct middle cerebral artery occlusion (MCAO) or sham operation for each *db/db* and *db/+* mouse, resulting in four groups: *db/+* sham, *db/+* MCAO, *db/db* sham and *db/db* MCAO. We used those mice of n=1 per group for scRNAseq and n=3-4 per group for bulk RNAseq.

### Experimental stroke

Focal cerebral ischemia was induced by distal MCAO ^25^ with modifications ^26,27^. The mice were initially anesthetized with 5% Isoflurane in a 30% O^2^ /70% N^2^O gas mixture for 30 to 60 seconds. Anesthesia was maintained by delivery of 2% Isoflurane during surgery via a facemask. The mice were placed on their sides and a 1-cm skin incision was created between the left margin of the orbit and the tragus. The masseter muscles were incised at the inferior edge of the zygoma, and then the mandible was pulled downward to expose the skull base. Using the dental drill, a small craniectomy was made above the proximal segment of the MCA, which could be seen through the exposure in the skull. The MCA segment just proximal to the olfactory branch was then transected with microscissors following electrocauterization to produce permanent occlusion. Just before the occlusion of the MCA, the left common carotid artery was temporally occluded with a microaneurysmal clip and reperfused 60 minutes later. Mice subjected to the same surgery without vessel occlusion served as sham-operated controls.

### Cell isolation

Mice were euthanized by anesthetic overdose using isoflurane three days after surgery. Peripheral blood was taken by heart puncture and mixed with acid-citrate dextrose solution (6:1, V: V, Santa Cruz Biotechnology) as previously described ^28^. In an effort to minimize purification time and preserve transcriptome, mononuclear cells were effectively isolated from the buffy coat from Ficoll-Paque density gradient (Catalog No. 17-5446-02, GE Healthcare Life Sciences, Chicago, IL) according to the manufacturer’s instructions. Briefly, for density gradient separation, 2 ml of blood and balanced salt solution mixture was added to 3 ml of Ficoll-Paque media (density: 1.084) and centrifuged at 500 × g for 30 min at 4°C without break. The mononuclear cell layers were recuperated and washed twice with PBS at 400 × g for 10 min each at 18°C.

### scRNA-seq

Single cells were encapsulated in droplets using 10X Genomics GemCode Technology and processed following manufacturer’s specifications. Briefly, every cell and every transcript are uniquely barcoded using a unique molecular identifier (UMI) and cDNA ready for sequencing on Illumina platforms is generated using the Single Cell 3’ Reagent Kits v2 (10X Genomics).

10X genomic single cell transcriptome sequencing data was processed using the Cell Ranger Single Cell software suite Version 1.3 as described previously ^29^. Briefly, the FASTQ files were processed with the Cell ranger count pipeline, which uses STAR^30^ to align the reads to mm10 mouse reference transcriptome. scRNAseq counts data and mean cluster transcript expression, as well as heatmaps of differentially expressed genes, were extracted with the Cell Ranger R kit. UMI normalization was done by dividing UMI counts by the total UMI counts in each cell, followed by multiplication with the median of the total UMI counts across cells. Feature counts from four samples were combined and analyzed with the R/Bioconductor package Seurat ^31^. Cells with less than 500 were filtered out; genes expressed in only 1 cell were omitted. Altogether, the filtered data contained 7808 cells. Expression data was log-normalized, and data were scaled to regress out differences in number of detected molecules. Variable genes were identified using the FindVariableGenes function with a x.low.cutoff = 0.0125, x.high.cutoff = 3 and y.cutoff = 0.5. For preliminary cell clustering, PCA was performed and appropriate components were visualized in two dimensions using uniform manifold approximation and projection (UMAP) ^32^.

We used SingleR ^33^ for the identification of various leukocyte populations including monocytes. The SingleR pipeline is based on correlating gene expression signatures of pure cell types published previously with gene expression data from single cells. We primarily used the ImmGen database as a reference data set, composing a collection of 830 microarray samples with 20 main cell types and further annotated to 253 subtypes ^34^. Clustering was performed using Ward’s hierarchical agglomerative clustering method based on the SingleR scores across all cell types ^33^ for the identification of monocyte population. Re-clustering focusing on specific cell types was performed for the monocyte population using Seurat.

### Fluorescence activated cell sorter (FACS)

We used flow cytometry for the validation of the diabetic effect on the cell population change in peripheral blood mononuclear cells. Blood was extracted in the same way as for scRNA-seq, and contaminating red blood cells were removed with RBC lysis buffer (Invitrogen) to retain granulocytes lost to the Ficoll gradient. Remaining cells were washed with PBS and blocked with 10% rat serum and then incubated with following fluorescent monoclonal antibodies from Invitrogen: CD11b eFluor450 (Clone M1/70), CD45 APC (Clone 30-F11), CD45R PerCP-Cyanine5.5 (RA3-6B2), Ly-6C APC-eFluor780 (HK1.4), TER-119 PE-Cyanine7 (TER-119), CD3 FITC (17A2), and Ly-6G PE-eFluor610 (1A8-Ly6G). Fixable viability dye eFluor506 was used to exclude dead cells. Cell staining was analyzed on a flow cytometer (BD FACSAria3) and further analysis was performed using FlowJo 10.0.

### Bulk RNAseq

Blood was extracted in the same way as for flow cytometry. Monocytes were isolated using EasySep^TM^ Mouse Monocyte Isolation Kit (STEMCELL Technologies) according to the manufacturer’s instructions. RNA was isolated by guanidine isothiocyanate/phenol purification followed by column cleanup. Briefly, the cerebellum was discarded, and each hemisphere was homogenized in QIAzol reagent (QIAGEN). The aqueous phase was separated by chloroform extraction (5:1, V: V), mixed with 70% ethanol (1:1), loaded on RNeasy Mini columns (QIAGEN) and processed according to the manufacturer’s guidelines. PolyA+ RNA-seq libraries were prepared from 500 ng total RNA input supplemented with 2 μl 1:200 ERCC RNA Spike-In Mix (Invitrogen) using the TruSeq Stranded mRNA HT Kit (Illumina) according to the manufacturer’s instructions.

Sequencing libraries were pooled and sequenced to a dept of > 7 million paired end 2×50 fragments on an Illumina HiSeq2500. After sequencing, raw RNA-Seq FASTQ files were generated by clustering BCL files using the Illumina Basespace BCL2FASTQ pipeline. RNA-seq analysis was performed in the commercial platform Omicsoft Arraystudio, including raw RNA-seq data quality control, filtering and trimming of reads, and alignment to the mouse genome (mm10 assembly, with gene annotations from Gencode release M6 and Ensembl81) using the Omicsoft OSA4 aligner 56, which is similar to the STAR aligner 57. The percentages of uniquely and nonuniquely mapped paired-end reads ranged from 86 to 92% and 3 to 6%, respectively, which was consistent with high-quality RNA-seq data. Gene-level quantification of aligned reads and differential gene expression analysis were performed using the Arraystudio implementations of RSEM and DESeq2, respectively. For differential gene expression analysis, a general linear model was used that included surgery (sham or MCAO) and T2DM genotype (*db/+* or *db/db*) as factors, and surgery: genotype as an interaction term. PCA and hierarchical clustering analysis to generate heatmaps was performed in Arraystudio.

### Bioinformatic analysis

We performed two pairwise comparisons for the differential gene expression analyses: T2DM vs. Control and T2DM+Stroke vs. Stroke. An adjusted p-value of <0.05 for scRNAseq and a false discovery rate (FDR) of <0.05 for bulk RNAseq served as cutoffs for significance between the groups. Significantly up- or down-regulated genes between groups were selected as the exclusively regulated genes and defined as DEGs in this study.

#### Pathway analysis for scRNAseq

Gene Set Enrichment Analysis (GSEA) if DEGs was performed using a collection of 50 refined “hallmark” gene sets derived from the Molecular Signature Database to convey specific and coherent biological state and minimize variation and redundancy ^35^. Complementary pathway enrichment using overrepresentation analysis was conducted using functional annotation tools DAVID (https://david.ncifcrf.gov/) ^36^. Significant GeneOntology (GO) terms of Biological Process (BP) and Kyoto Encyclopedia of Genes and Genomes (KEGG) pathways were identified with adjusted p-value of <0.05 set as the cutoff criterion. We additionally assessed the association between DEGs and genes previously reported to be related to neuroprotection in myeloid cells ^18,37^.

#### Pathway analysis for bulk RNAseq

In addition to GSEA mentioned above, we also performed overrepresentation analysis using Metascape ^38^. Genes differentially expressed were identified from bulk RNA sequencing data, selecting those with a log2 fold change greater than 1 and an adjusted p-value less than 0.05. For each curated list of genes, comprehensive pathway and enrichment analyses were conducted utilizing several ontology sources, including GO BP, KEGG Pathway, Reactome Gene Sets, and Wiki Pathways. Significant terms were identified based on a p-value threshold of less than 0.01, a minimum occurrence count of three, and an enrichment factor exceeding 1.5, which represents the ratio of observed counts to counts expected by chance. These terms were then aggregated and organized into clusters according to their similarity in membership.

### Lipopolysaccharide (LPS) preconditioning

We compared the effect of preconditioning via Toll-like receptor 4 agonist LPS on neuroprotection against ischemic stroke between *db/+* and *db/db* mice. Ultrapure LPS from the Salmonella Minnesota R595 strain was obtained from InvivoGen. Mice (*db/+* and *db/db*) were injected with either purified LPS (0.2 mg/kg, s.c.) or saline (100 µl, i.p.) three days before MCAO. Neurological assessment was performed 24 hrs after MCAO using a 24-point scoring system comprised of 4 sensory and 4 motor assessments as previously reported ^39,40^. Mice were euthanized followed by transcardiac perfusion with 4% PFA 3 days after MCAO for lesion assessment. Infarct volume was determined via TTC-stained sections by Fiji ^41^. Percentage of infarction was calculated by dividing infarct volume with total brain volume.

### Western Blot Analysis

Three days after MCAO, LPS or sham, ipsilateral and contralateral sides of brain were dissected and homogenized in RIPA buffer (radioimmunoprecipitation assay buffer) (1% Triton-X100, 0.5% sodium deoxycholate, 0.1% SDS, 25mM Tris, 150mM NaCl, pH 7.4) with 1X protease and phosphatase inhibitor cocktail (Cat#78429 and 78420; Thermo Fisher Scientific) immediately. The proteins from cytoplasm and nucleus were collected separately. The protein concentration was determined using Pierce^TM^ BCA protein assay kits (Cat#23225; Thermo Fisher Scientific). Brain extracts containing 60 ug of protein were loaded and separated on 6-10% denaturing polyacrylamide gel electrophoresis (SDS-PAGE) and transferred to polyvinylidene difluoride (PVDF) membrane (Cat#88518; Thermo Fisher Scientific). The membrane was blocked with 5% milk in Tris-buffered saline with 0.1% Tween (TBST) and incubated overnight with primary antibodies: mouse anti-STAT1 (1:800, Cat# 603702, Biolegend), mouse anti-STAT1 phospho (Ser727) (1:800, Cat#686402, Biolegend), mouse anti-IRF3 ((1:1000, Cat# 655702, Biolegend), rat anti-IRF7 ((1:500, Cat#696402, Biolegend), rat anti-IRF8 ((1:500, Cat# 623102, Biolegend) at 4 °C, followed by goat anti-mouse and anti-rat IgG (H+L) conjugated to horseradish peroxidase (HRP) (1:4000, Jackson Immunochemicals) for 1 h at room temperature. After washing, the immune complexes were detected by the SuperSignal West Pico PLUS Chemiluminescent Substrate (Cat # PI34577, Fisher scientific). β-actin (1:5000, Cat # 20536-1-AP, and 66009-1-Ig, Proteintech) was used for normalization for tissue samples ^42^.

### Meningeal gene expression analysis

Three days after LPS injection, meninges were harvested from the calvaria as previously described ^43^. In brief, the meninges were scraped from the skullcap in 1 x PBS and were homogenized in TRIzol (Cat#15596018, Life Technologies). Following homogenization, the RNA was extracted using phenol-chloroform method and then cleaned up with RNA Mini Kit (Qiagen). The quality and concentration of the RNA was assessed with Agilent RNA 6000 Pico Kit (Agilent, Cat#5067-1513) on Bioanalyser 2100 (Agilent). Gene expression was determined using Nanostring nCounter® Myeloid Innate Immunity Panel v2. Data normalization and quantification were performed using the nSolver 4.0 software (NanoString, Seattle, WA, USA) as recommended by the manufacturer. Following normalization, to compare the effects of LPS, we calculated the fold change of genes involved in the IFN pathway after LPS stimulation relative to the unstimulated level for both db/+ and db/db mice. Three biological replicates were obtained for each group.

### Statistical analysis

Values are shown as means ± SD. For the analysis of flow cytometry, we used paired two-tailed t-Tests. For other results, we used two-way ANOVA followed by post-hoc pair-wise comparisons and p-values adjusted for multiple comparison. A p-value of <0.05 was considered statistically significant. Data were analyzed and graphs created using GraphPad Prism 10 software.

## Results

### Single-cell RNA sequencing identifies a T2DM-specific composition of murine peripheral blood mononuclear cells

To gain insights into how T2DM and stroke affect the landscape of peripheral immune cells with respect to cellular composition and transcriptome, we performed scRNAseq from peripheral blood mononuclear cells isolated 3 days after stroke or sham surgery in diabetic db/db mice and normoglycemic control db/+ mice. Following the application of quality control filters, a total of 7808 cells were included in the analysis. Via unsupervised clustering analysis, 16 distinct cell clusters emerged. The cell types were identified using SingleR and main cell types among clusters were used for annotation (Fig. 1*A*). Monocytes formed the biggest population compared to other cell types such as T cells, B cells, NK cells, dendritic cells, and neutrophils. We confirmed the annotation of each cell type by mapping the gene expression patterns of established canonical markers on UMAP plots (Fig. 1*C*). The relative percentage of monocytes increased, and lymphocytes reduced comparing *db/db* to *db/+* groups but remained similar comparing stroke to non-stroke (Fig. 1*B*, *D*). Flow cytometry data confirmed a similar change in the percentages of monocytes and lymphocytes in *db/db* compared to *db/+* non-stroke (Fig. 1*E, F*). In addition to the increase in monocytes, we found that there was also a significant increase of neutrophils in *db/db* compared to *db/+* non-stroke mice (Fig. 1*E, F*).

**Figure 1.**
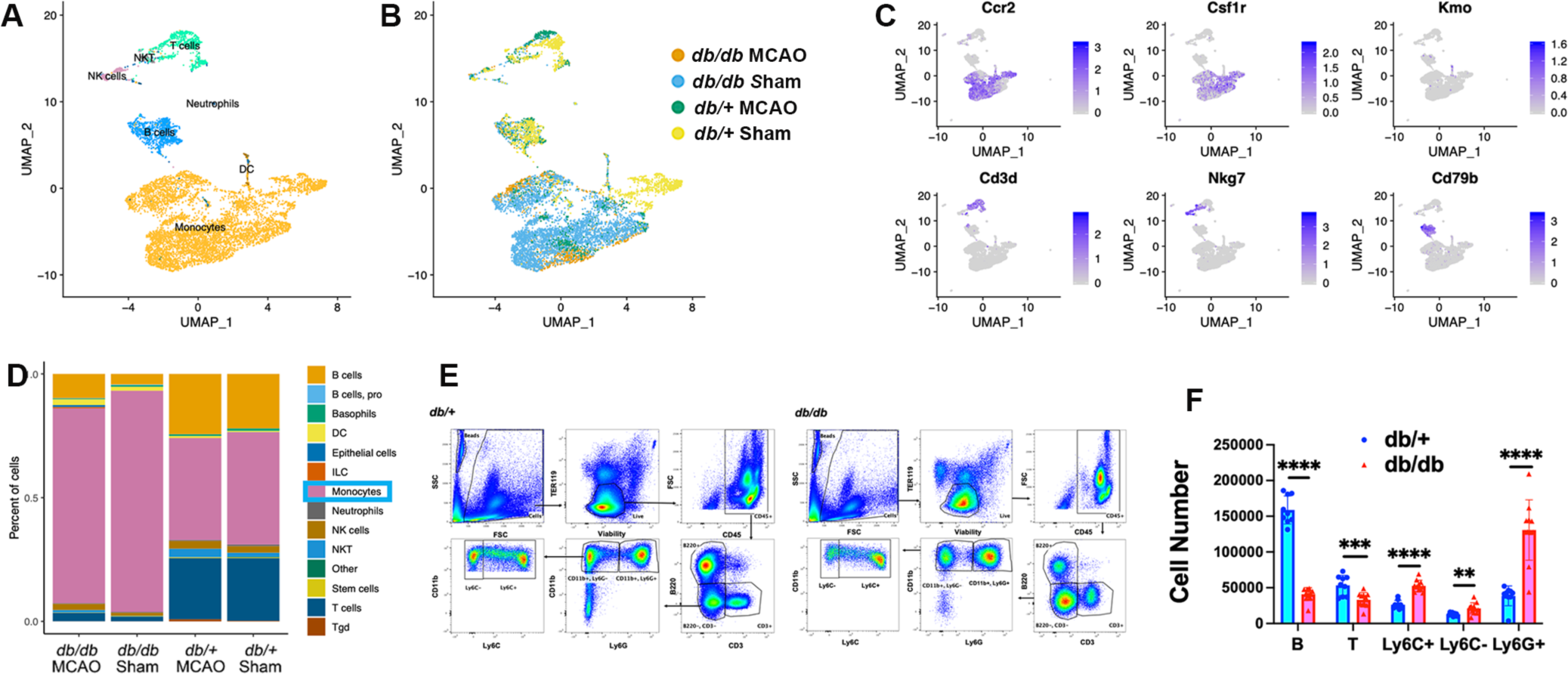
Single-cell RNA sequencing analysis of peripheral mononuclear cells and experimental validation using flow cytometry. *A, B*, the uniform manifold approximation and projection (UMAP) plot of aligned gene expression data in single cells extracted from 4 experimental groups of peripheral mononuclear cells showing 16 distinct clusters with 7 major cell types (*A*) and showing experimental group (*B*). *C*, Gene expression patterns of established canonical markers projected onto UMAP plots (scale: log-transformed gene expression). Markers are selected to distinguish monocytes from other leukocyte cell types: monocytes (*Ccr2* and *Csf1r*), dendritic cells (*Kmo*), T cells (*Cd3d* and *Nkg7*), and B cells (*Cd79b*). *D*, stacked bar plots showing the proportions of each cell type in each group of mice. *E*, Gating strategy used to identify immune cell subsets in FACS analysis of PBMCs from *db/+* and *db/db* mice. After gating based on size and excluding dead cells and red blood cells, live leukocytes are defined as CD45+ population. B cells, T cells, and myeloid cells are defined as CD45R(+), CD3(+), and double negatives, respectively. Monocytes and neutrophils are distinguished among the myeloid cells using CD11b and Ly6G, and monocytes are divided into Ly6C- and Ly6C+ populations.: lower row. *F,* Scatter plots comparing the number of major cell types in PBMCs following RBC lysis between *db/+* and *db/db*. *db/db* mice using flow cytometry shows significantly increased proportions of Ly6C+ monocytes and neutrophils and decreased proportion of B cells compared to control *db/+* mice (n = 5 each, \*\**p*< 0.05, \*\*\**p*< 0.01, \*\*\*\**p*<0.005). Data are shown as means ± SD.

### Identification of monocyte subsets in normal and disease conditions

#### Re-clustering for monocytes

A total of 5761 myeloid cells annotated by SingleR were included in the initial Seurat-based unsupervised re-clustering, resulting in 8 distinct clusters (Fig. 2*A*). Cluster 4 is almost exclusively monocytes from the db/+ sham group (Fig. 2*B*, *C*). Conversely, other clusters were proportionally enriched with the three pathological groups relative to the db/+ sham group. Overall, the relative proportions of Ly6C+ and Ly6C-populations were comparable among 4 experimental groups (Fig. 2*D*).

**Figure 2.**
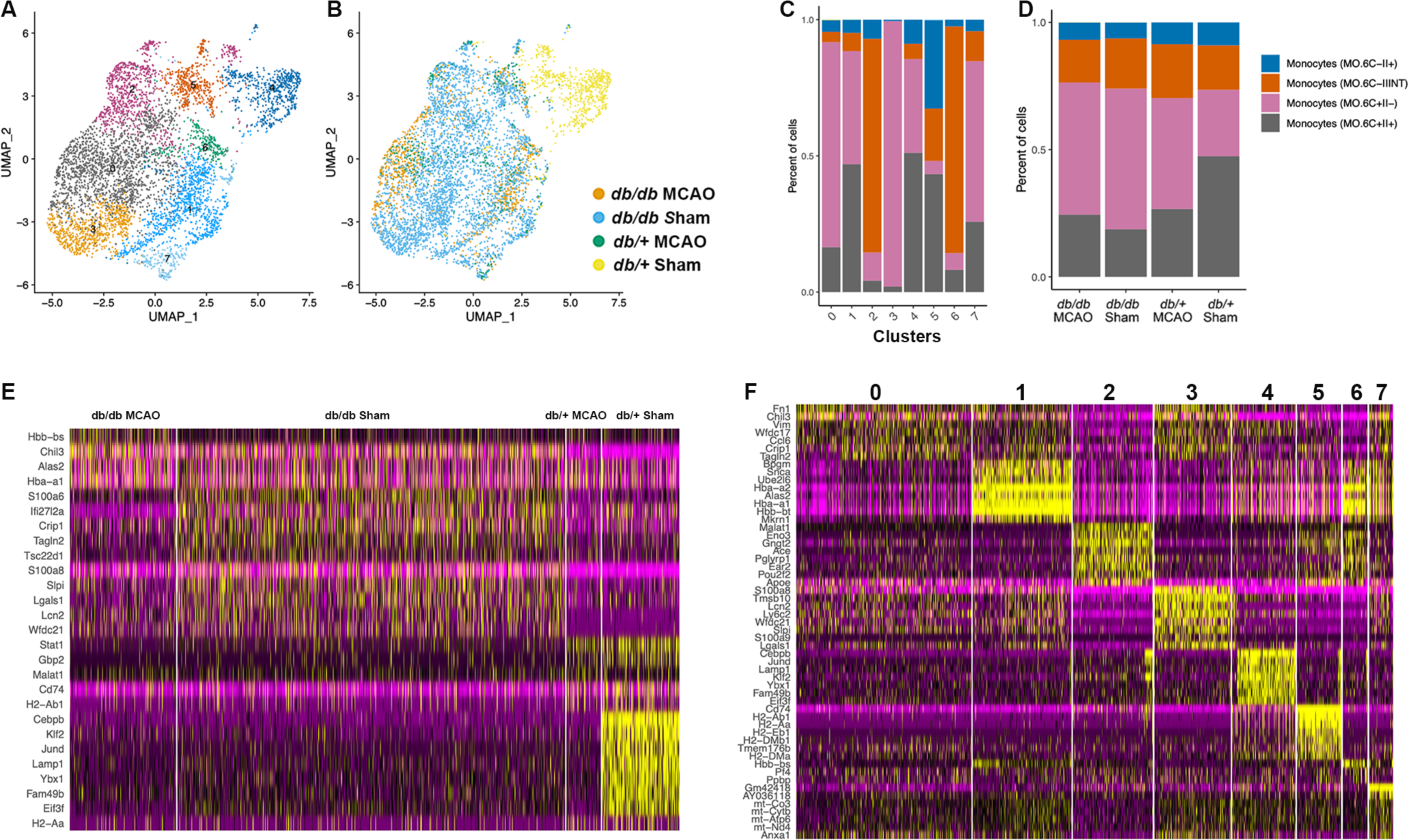
Re-clustering of myeloid population from single-cell RNA sequencing data. *A, B*, the uniform manifold approximation and projection (UMAP) plots of aligned gene expression data in single cells extracted from all 4 experimental groups of monocytes showing 8 distinct clusters (*A*) corresponding to 4 experimental groups (*B*). *C&D*, stacked bar plots showing the proportions of each monocyte subtype annotated by SingleR in each cluster (C) and experimental group of mice (D). *E&F*, Heatmap showing the most upregulated genes (ordered by decreasing p-value) uniquely in each cluster (*E,* top 5 genes) and experimental group (*F,* top 8 genes). Most notably cluster 4 is almost exclusively comprised of cells from the db/+ sham group that lacks the influence of stroke or type 2 diabetes, and expressed elevated levels of *Cebpb, Klf2, Jund, Lamp1, Ybx1, Fam49b* and *Elf3f* genes compared to any other group. Scale: Log2 fold change.

Per monocyte classification scheme established by ImmGen, clusters 2, 5 and 6 contain the least proportion of MHCII− monocytes and exhibit higher levels of expression for *Eno3, Gngt2, Ace*, and *Apoe* genes than other clusters. Clusters 2 and 6 consist mainly Ly6C-monocytes expressing intermediate levels of MHCII, characteristic of patrolling monocytes. Although cluster 5 has comparable proportions of Ly6C+ and Ly6C-cells, it exhibits strong expression of MHCII genes such as *H2-Ab1, H2-Aa*, and *Cd74*, consistent with having the lowest percentage of MHCII− monocytes among all clusters ^44,45^. In contrast, clusters 0, 3 and 7 are dominated by Ly6C+ monocytes characteristic of the classical inflammatory monocytes. Cluster 3 is almost exclusively represented by the db/db non-stroke group and almost entirely Ly6C+MHCII−, exhibiting strong expression of pro-inflammatory genes such as *Ly6c2, Lgals1, S100a8*, and *Lcn2* (Fig. 2*E*) and representing a subset recently described as neutrophil-like monocytes ^46–48^. Clusters 1 and 6 showed elevated expression of genes related to heme metabolism such as *Hba, Hbb, Alas2, Snca,* and *Bpgm*, with particularly high expression of hemoglobin subunit *Hba* and *Hbb* genes. Among these, the hemoglobin beta-subunit adult s chain (*Hbb-bs*) was specifically increased in db/db mice with stroke (Fig 2E&F). Cluster 4, which had strong expression of *Cebpb, Jund*, and *Klf2* but low expression of *S100a8* and *Lcn2*, was a major Ly6C+ subset in db/+ non-stroke group, suggesting that this subset is equivalent to the classical monocytes at steady state (Fig. 2E, F).

### Prominent effects of diabetes and stroke on peripheral monocytes transcriptome

Monocytes in diabetic stroke and non-stroke mice exhibited a greater heterogeneity in transcriptome patterns compared to non-diabetic mice (Fig. 2*F*). Compared to the db/+ sham or control group, pro-survival genes such as *Cebpb, Klf2, Jund, Lamp1, Ybx1, Fam49b*, and *Eif3f* were downregulated in all the other pathological groups affected by T2DM and/or stroke. T2DM upregulated the expression of inflammatory genes such as *S100a8, Chil3, Lcn2*, as well as genes involved in metabolic or growth promoting changes like *Wfdc21* (myeloid-derived suppressor cells: promote tumor metastasis), *Slpi* (Secretory leukocyte protease inhibitor: anti-inflammatory or growth-promoting) and *Crip1* (oncogene). In contrast, T2DM downregulated genes involved in interferon signaling including Signal Transducer and Activator of Transcription 1 (*STAT1*) (both type I and type II IFN) and guanylate binding protein 2 (*Gbp2*) (cellular response to interferon-beta), as well as genes relevant to antigen presentation such as *Cd74* and *H2-Ab1*, suggesting that T2DM may lead to immunosuppression.

### Specific biological pathways affected by diabetes and stroke

Gene Set Enrichment Analysis using available 50 hallmark pathways ^35^ found that in both Ly6C+ and Ly6C-monocytes, pathways involving IFNα- and IFNγ-response were strongly activated in db/+ mice with and without stroke, but only very weakly activated in the db/db stroke mice. IL6_JAK_STAT3 signaling was weakly activated in db/+ mice, but negative in db/db monocytes. Pathway involving allograft rejection is almost only acted in the db/+ mice, while pathway in heme metabolism is uniquely upregulated in T2DM mice with stroke (Fig. 3). To confirm the transcriptome changes and biological pathways detected by scRNAseq in monocyte populations caused by diabetes or stroke, we carried out bulk RNAseq analysis of the isolated peripheral blood monocytes from the same experimental groups, of which the heatmaps are shown in Fig 4A. In both scRNAseq and bulk RNAseq analyses, T2DM groups displayed down-regulated pathways related to type 1 or type 2 IFN signaling (Table S1, Fig S1). Apart from many shared down-regulated genes in IFN signaling and antigen presentation in the T2DM groups between scRNAseq and bulk RNAseq, the latter also detected downregulation of IL27 signaling in T2DM groups based on overrepresentation analysis, leading to reduced activation of receptor-associated Janus Kinases and nuclear translocation of STAT1 and STAT3 transcription factors (Fig 4B). Further, bulk RNAseq identified additional T2DM upregulated pathways in inflammatory response such as leukocyte migration and neutrophil degranulation, complement and coagulation, as well as those involved in complement receptor signaling such as Serpinb2, VWF, coagulation factors II/V/VII, C1q, C3ar1, C5ar1, and Cd59 (Fig 4C). Not surprisingly, the T2DM upregulated pathways in inflammation and coagulation are more pronounced in the stroke compared to sham groups. Top DEGs involved in the IFN, antigen presentation and complement/coagulation pathways comparing db/db to db/+ monocytes in non-stroke and stroke conditions are listed in Tables S2, S3 and S4 Interestingly, several pathways related to leukocyte migration and neutrophil degranulation were commonly upregulated between T2DM and stroke, implying the potential of T2DM in exacerbating stroke-induced inflammation and immune cell chemotaxis (Fig 4D). Lastly, pathway in cholesterol biosynthesis was downregulated by stroke in monocytes of both diabetic and non-diabetic mice (Fig 4E).

**Figure 3.**
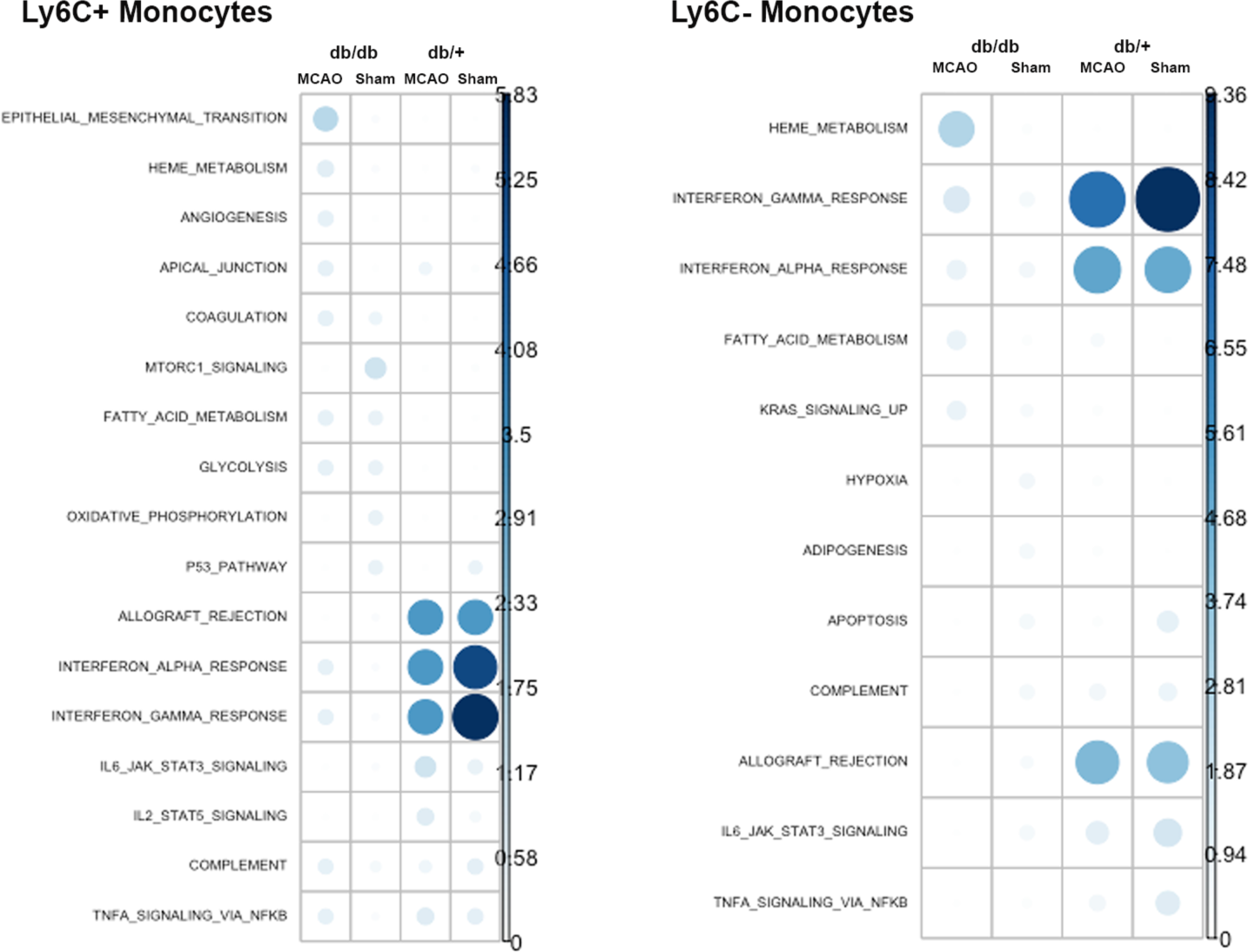
Gene ontology (GO) enrichment analysis revealed downregulated signaling in type 1 and type 2 interferon and allograft rejection signaling pathways in db/db monocytes compared to those of db/+ under non-stroke and stroke conditions. T2DM effect on pathways are similar between Ly6C+ and Ly6C-monocytes. Scale for intensity based on −log10 (adjusted p-value) for enrichment of marker genes per group in 50 hallmark pathways (capped by 10 which is p val=1e-10).

**Figure 4.**
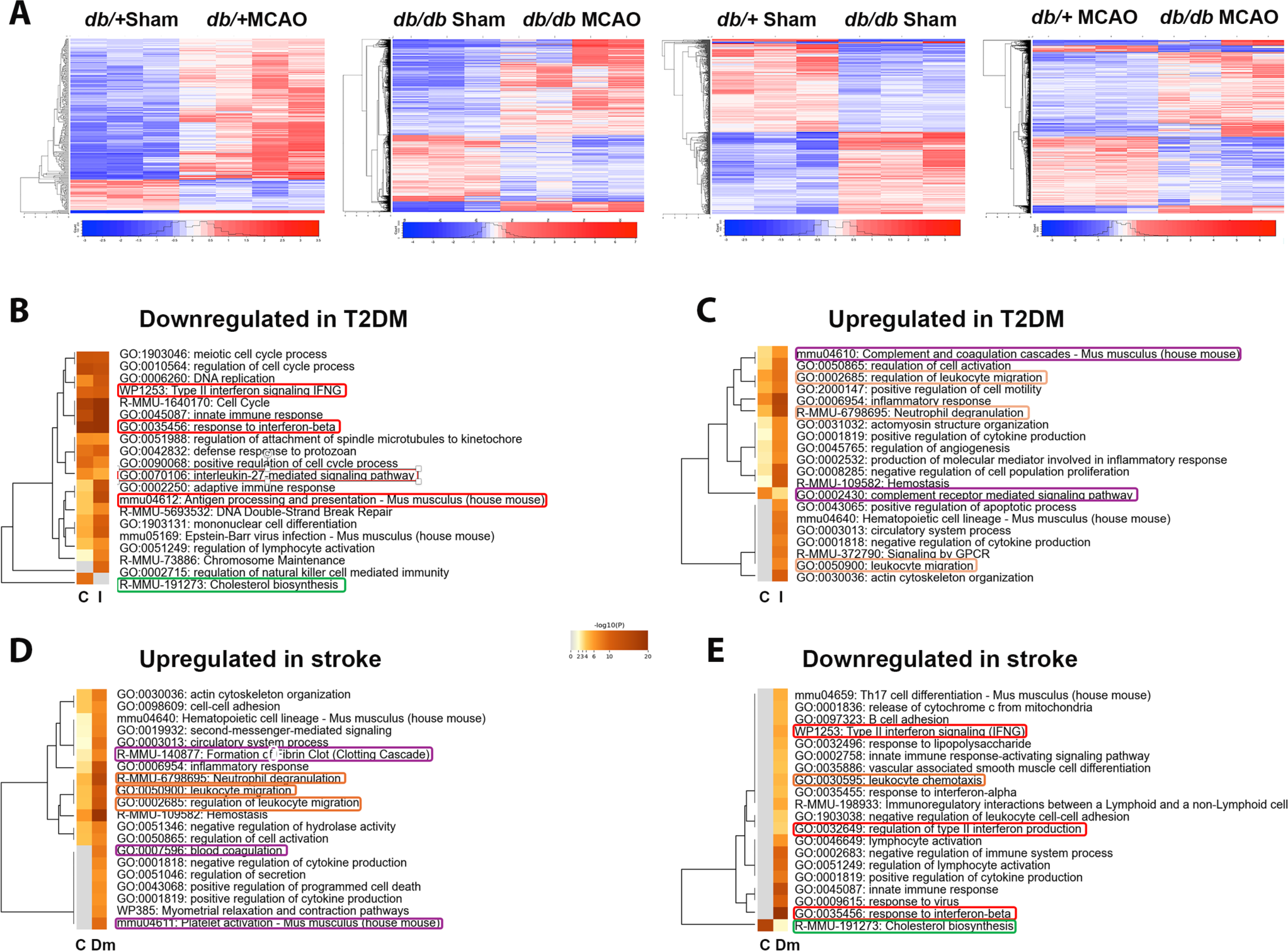
Bulk RNA sequencing revealed downregulated signaling in IFN pathways and upregulated pathways involving coagulation, complement and chemotaxis in T2DM monocytes compared to control monocytes. *A*, Heatmaps of gene expression in 4 pair-wise comparisons between db/+ Sham and db/+ MCAO, db/db Sham and db/db MCAO, db/+ Sham and db/db Sham, and db/+ MCAO and db/db MCAO, arranged by hierarchical clustering for genes and RNA samples to reveal maximal differences in patterns between groups. N=3 biological replicates/sham group, N=4 biological replicates/MCAO group. *B-E*, Gene ontology (GO) enrichment analysis of biological processes was performed with filtered DE genes comparing T2DM or stroke effect. DEGs were filtered when log2Fold change > 1 and FDR <0.05. Significance of enrichment in log10(p) is shown in the color shades. Among others, pathways in type 1- and type 2-IFN signaling, IL27-mediated signaling pathway and those regulate antigen processing and presentation were downregulated in T2DM monocytes in both stroke (I) and non-stroke (C) conditions. Pathways in complement and coagulation cascades and complement receptor signaling were upregulated, together with leukocyte migration in both non-stroke and stroke conditions. Formation of fibrin clot and leukocyte migration pathways were upregulated in both non-diabetic (C) and diabetic (Dm) monocytes, although blood coagulation and platelet activation pathways were only upregulated in the latter. Cholesterol synthesis pathway was downregulated in both control and diabetic monocytes, while IFN pathways were downregulated only in the latter.

Moreover, many interferon-stimulating genes (ISGs) previously reported to be neuroprotective in the brain ^18,37^ were downregulated in the monocytes of db/db mice compared to db/+ mice after stroke (Table 1). To determine the effect of stroke and T2DM on IFN signaling in the brain, we quantified the expression of several key signaling molecules in the IFN pathways in brain homogenates by western blot analysis. Two-way ANOVA suggests that there was a strong stroke effect on the induction of IRF3 (p<0.0001) and IRF7 (p<0.0001) comparing ipsilateral to contralateral sides and a significant diabetes effect on the expression of pSTAT1 (p<0.001) and IRF3 (p<0.05). Specifically, stroke-induced Phosphorylated-Signal-Transducer-And-Activator-Of-Transcription-1 (pSTAT1) as well as IRF3 were reduced in the brain homogenates ipsilateral to stroke of db/db mice compared to db/+ mice 3 days after dMCAO, consistent with the T2DM associated suppression in IFN signaling detected in monocyte transcriptome data (Fig 5).

**Figure 5.**
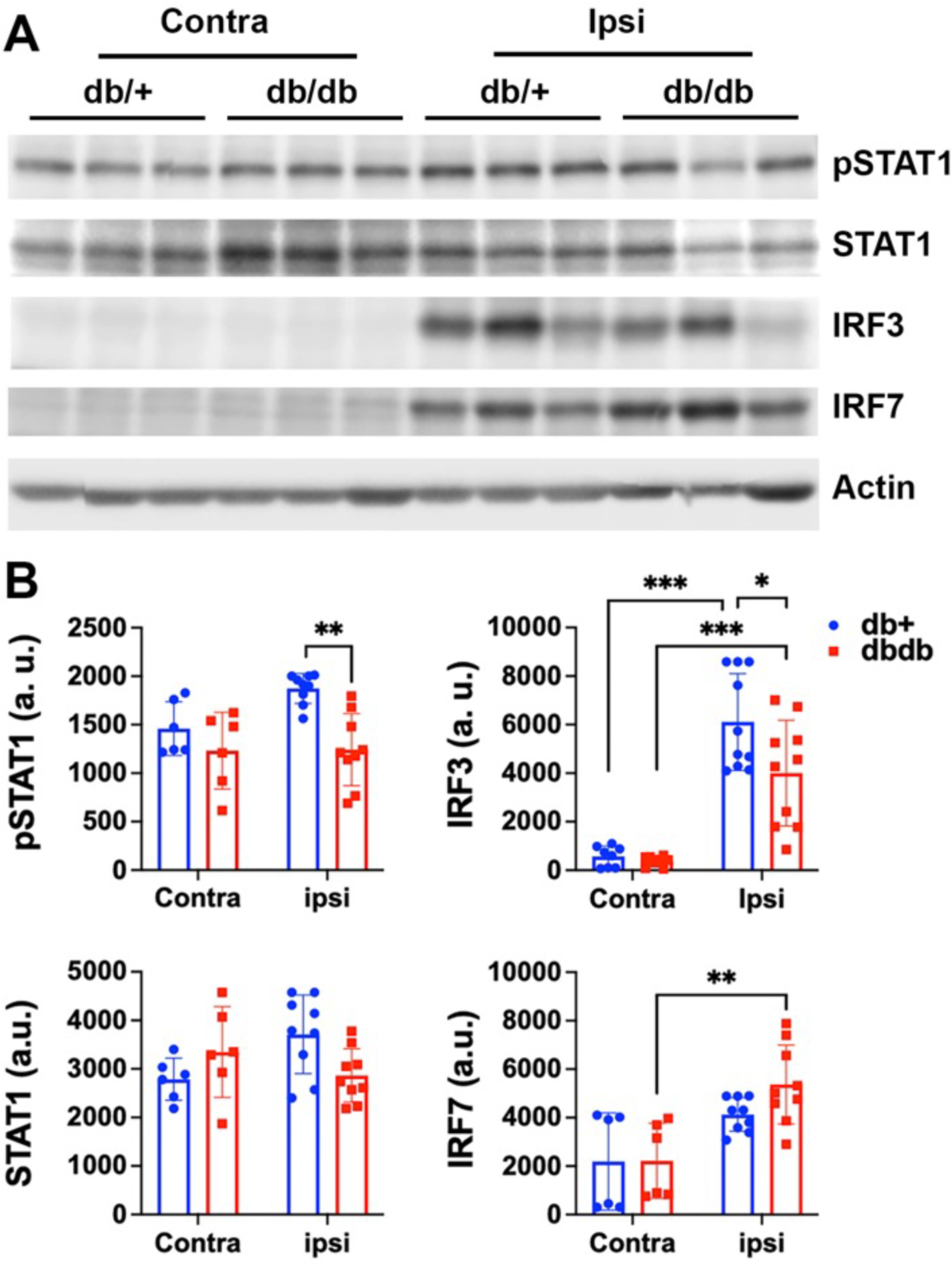
T2DM reduced stroke-activated interferon signaling in the brain. Three days after dMCAO, brain homogenates from ipsilateral ischemic regions and homotopical contralateral regions were analyzed by western blots for tyrosine phosphorylated-STAT1 (pSTAT1), STAT1, IRF3, IRF7, and beta-Actin. *A*, representative blots for the above markers. *B*, the intensity of the western blot band for the IFN markers was quantified and shown in the graphs. Two-way ANOVA detected a significant genotype effect for pSTAT1 (p<0.001) and IRF3 (p<0.05), as well as a significant stroke effect for IRF3 (p<0.001) and IRF7 (p<0.0001) by comparing ipsilateral to contralateral side. There were significantly less pSTAT1 and IRF3 in the ipsilateral brain homogenates from db/db mice compared to db/+ mice after stroke. Contra: N=6/maker/group; Ipsi: 9-10/marker/group.

**Table 1.**
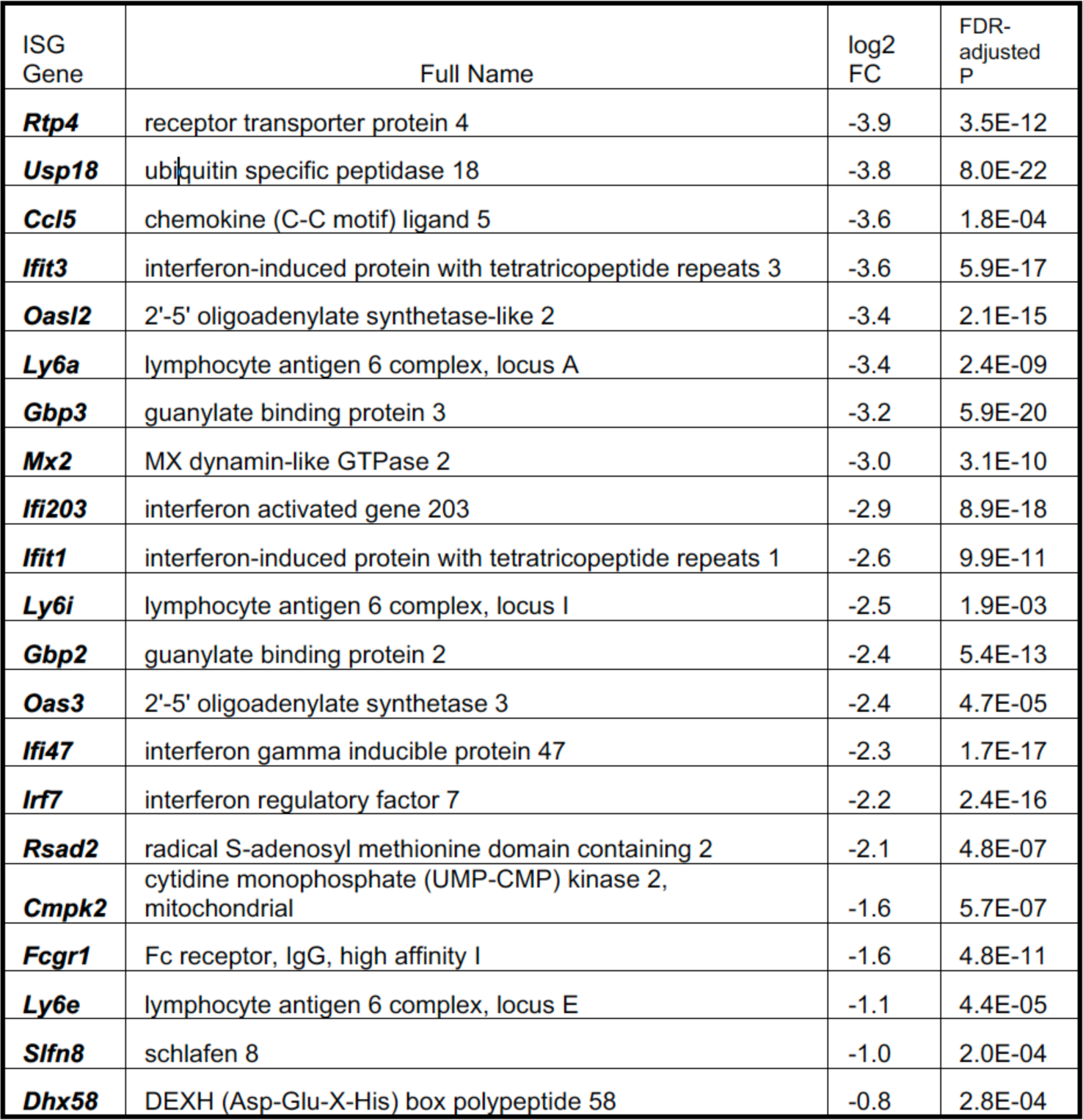
T2DM reduced the expression of interferon stimulated genes (ISGs) in monocytes after ischemic stroke. Examples of differentially expressed interferon stimulated genes (ISGs) in the peripheral monocytes at 3 ds after dMCAO determined by bulk RNAseq. Negative log2FC representing the fold change reduction in log2 scale comparing the db/db MCAO to *db/+* MCAO groups are shown for each listed gene with adjusted false detection rate (FDR) akin to P values. N= 3-4/group.

### LPS preconditioning failed to confer neuroprotection against stroke injury in diabetic mice

Preconditioning induced neuroprotection via brief ischemia or Toll-Like Receptor (TLR) agonists converges on interferon regulatory factor (IRF)-dependent signaling ^18,19,23^. Given that the IFN response is defective in the db/db mice, we hypothesize that the diabetic mice would show attenuated tolerance to stroke following Toll-like receptor-mediated preconditioning compared to normoglycemic db/+ mice. To test this hypothesis, we induced ischemic stroke 3 days after LPS administration. Modified neurological severity score (mNSS) was determined 1 day after MCAO via a 24-point composite scoring system and lesion sizes were assessed by TTC staining 3 days after MCAO. We found that db/+ mice had smaller stroke volume and higher mNSS score compared to db/db mice in both vehicle and LPS injected groups (Two-Way ANOVA, T2DM effect, infarct: P<0.0001; mNSS: p<0.0001). There is an overall LPS preconditioning effect in infarct reduction but not on neurological score (Two-Way ANOVA, PC effect, infarct: P<0.005). However, LPS PC only significantly reduced infarct size and increased neurological scores in db/+ but not in db/db mice (Fig 6A, B), consistent with the downregulated IFN signaling detected in T2DM mice.

**Figure 6.**
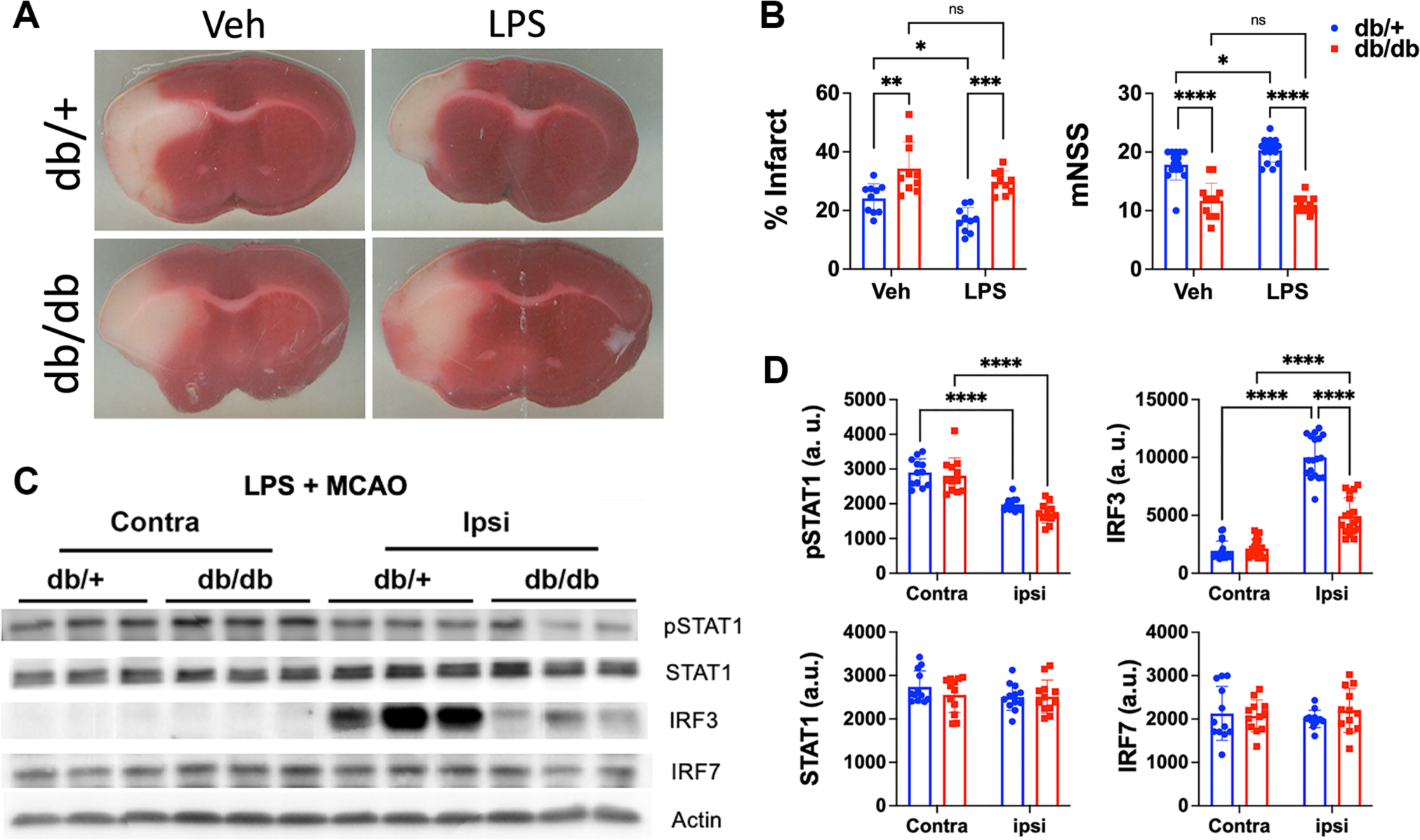
LPS preconditioning induced neuroprotection in db/+ but not in db/db mice, coincided with an increased induction of IRF3 in the ipsilateral brain. Three days after LPS injection, stroke was induced by dMCAO. One day after stroke, mNSS were determined. Three days after stroke, infarct size was determined and brain homogenates from ipsilateral ischemic regions and homotopical contralateral regions were analyzed by western blots. *A*, Representative TTC stained brain slices collected 3 days after MCAO surgery from groups as indicated. *B*, Quantification of the % infarct volume of 3 days and neurological scores (mNSS) determined one day after MCAO based on a 24-point composite score system. db/+ mice had smaller stroke volume and higher mNSS scores compared to db/db mice in both vehicle and LPS injected groups. LPS preconditioning significantly reduced infarct volume and improved mNSS in db/+ mice but did not in the db/db mice. *C*, Representative blots for tyrosine phosphorylated-STAT1 (pSTAT1), STAT1, IRF3, IRF7, and beta-Actin. *D*, the intensity of the western blot band for the IFN markers was quantified and shown in the graphs. Data were analyzed by two-way ANOVA and Bonferroni post hoc test adjusted for multiple comparisons. We found that the expression of IRF3 was significantly reduced in the db/db brains compared to db/+ brains that underwent LPS PC followed by stroke (p<0.0001), while pSTAT1 levels were reduced in ipsilateral compared to contralateral brain homogenates in both genotypes. *p<0.05; **p<0.01; ***p<0.005; ****p<0.001. Infarct: N=10/group. mNSS:11-15/group. Western blot: Contra: N=12/marker/group; Ipsi: 12/marker/group.

### Effect of stroke and LPS preconditioning on IFN signaling

We determined the effect of LPS PC on IFN signaling after stroke by western blot analysis, and we found that there was a significant effect of LPS PC followed by stroke on the induction of IRF3 (Two-Way ANOVA, hemisphere effect: p<0.0001) and decrease in pSTAT1 (hemisphere effect: p< 0.0001) expression in brain homogenates comparing ipsilateral to contralateral hemispheres (Fig 6C, D). Specifically, ipsilateral IRF3 level was significantly reduced in db/db homogenates compared to db/+ ones (p<0.0001), consistent with the lack of neuroprotection observed in db/db mice following LPS PC (Fig 6C, D). Since meninges was identified as a key immunoregulatory compartment involved in LPS PC-mediated neuroprotection ^16^, we also determined whether LPS altered the expression of genes in the IFN pathway in the meninges using total RNA probed with a panel of mouse myeloid markers. We found that there was a downward trend in the expression of genes in the IFN pathways in the meninges of db/db mice compared to db/+ mice one day after LPS treatment, with irf7 being significantly reduced, further linking the lack of preconditioning-induced neuroprotection to IFN signaling defect (Fig 8).

**Figure 7.**
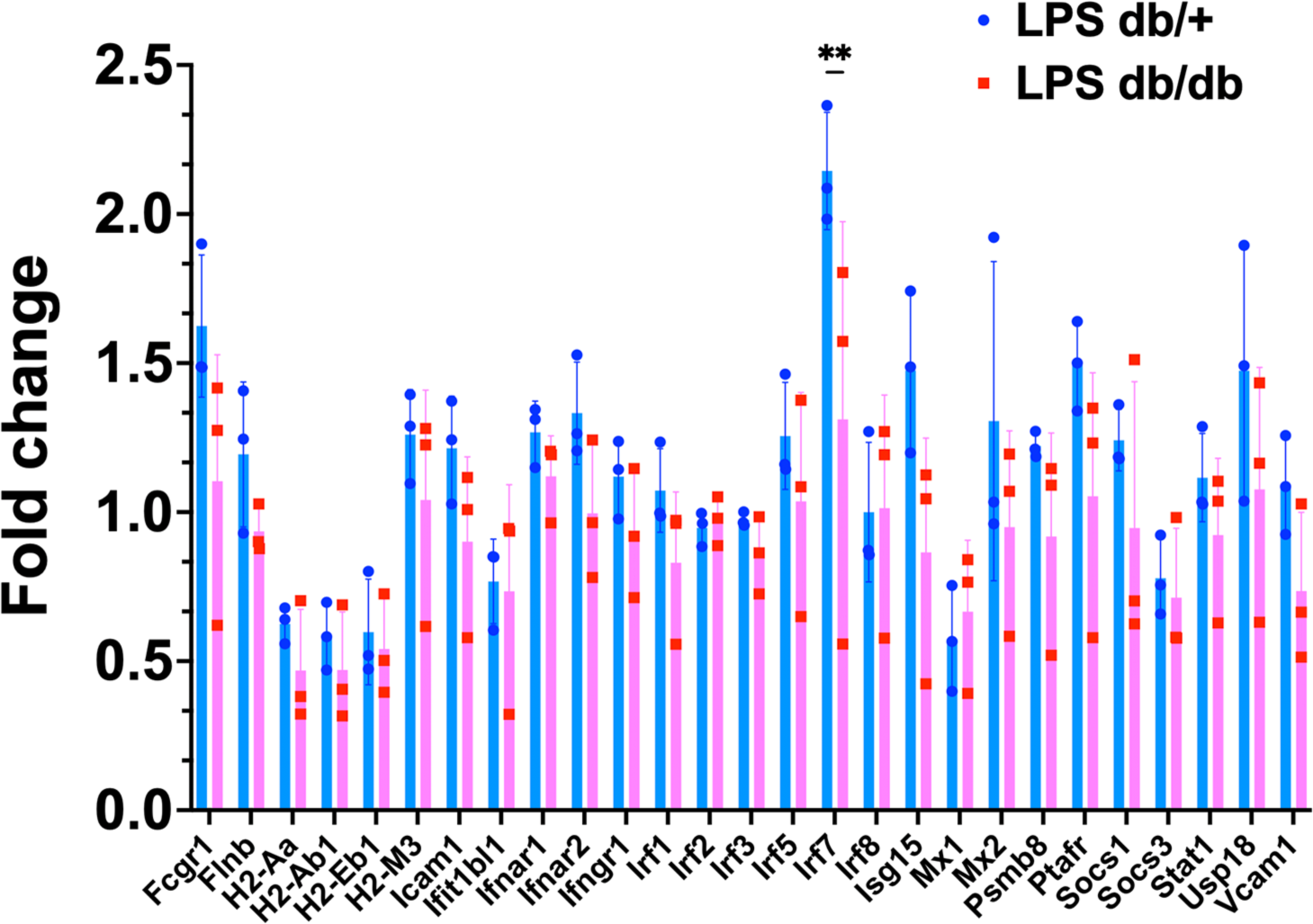
T2DM significantly reduced the expression of irf7 in the meninges after LPS stimulation. Total RNA was extracted from the meninges 3 days after LPS injection (0.25 mg/kg, i.p.) and detected using Nanostring nCounter platform and analyzed with nSolver. The expression of irf7 was significantly reduced in the meninges of db/db mice compared to db/+ mice, although there was a trending reduction in the expression of almost all the genes involved in the interferon signaling among the panel of probes. It suggests that LPS failed to activate the IFN pathways in the diabetic mice as robustly as in the control mice (Biological replicates N=3/group).

## Discussion

While the link between T2DM pathophysiology and chronic inflammation is well established, the effect of metabolic disturbance on immune cell function and potential impact on stroke outcome is less understood. Using scRNAseq and bulk RNAseq, our study revealed distinct changes in cell composition of peripheral blood immune cells and the monocyte transcriptome under the conditions of T2DM and stroke. The main findings of our study are that T2DM disproportionally increased myeloid cells but reduced lymphocytes in the peripheral blood, and that T2DM increased the expression of proinflammatory genes and genes involved in coagulation, accompanied by reduced expression of genes in the IFN pathways and genes related to antigen presentation. The defect in IFN signaling led to a dampened response to TLR4 receptor-mediated preconditioning and neuroprotection against ischemic injury. To our knowledge, this is the first report that revealed a major pathophysiological mechanism of how T2DM may exacerbate stroke inflammation and outcome combining scRNAseq and bulk RNAseq approaches.

### Monocyte diversity in diabetes and stroke

Recent advances and feasibility in scRNAseq enabled the unbiased mapping of cellular heterogeneity, uncovered highly variable genes contributing to the heterogeneity, characterized previously unrecognized disease-associated cell populations, and identified potential molecular regulators and pathway-specific markers ^49,50^. Emerging studies using this technology have acknowledged that monocytes are much more heterogeneous than previously recognized, revealing the existence of distinct subsets within Ly6C+ and Ly6C-monocytes based on the level of class II MHC gene expression. In general, Ly6C-non-classical monocytes display positive or intermediate levels of MHCII, while Ly6C+ classical monocytes are divided into two subgroups with either positive or negative MHCII expression. Our study has found that the T2DM groups have overall greater proportions of Ly6C+ monocytes negative for MHCII gene expression, while the control non-stroke group has the biggest proportion of monocytes positive for MHCII gene expression, regardless of Ly6C phenotype. It suggests that T2DM rendered the circulating monocytes with reduced potential for antigen presentation, as evidenced by the downregulated H2 gene expression found by both RNAseq studies. Ly6C-IIint monocytes expressing an intermediate level of MHCII genes were reported to be the transitional state before differentiating into Ly6C-monocytes from Ly6C^hi^ precursors egressed from bone marrow to circulation ^44,45^ ^44^. Interestingly, monocytes in the non-pathological control group consist of predominately Ly6C+ positive or negative for MHCII expression, accompanied by small populations of Ly6C-subsets. On the other hand, there was an increase of neutrophil-like monocytes in the T2DM groups, which represent a minor pool of Ly6C+ monocyte subsets maintaining homeostasis under steady-state, yet become conspicuous during inflammation ^46^. This subset is characterized by the elevated expression of genes related to granule proteins that are highly expressed by neutrophils such as S100a8 and Lcn2, accompanied by reduced expression of MHCII genes. Neutrophil-like monocytes are differentiated from granulocyte-monocyte progenitors (GMPs) and are functionally distinct from their Ly6C+ counterparts derived from monocyte-dendritic cell progenitors (MDP) ^46–48^. It is believed that classical monocytes derived from GMP and MDP pathways are functionally related to their neutrophil and dendric cell cousins, respectively ^46^. However, how T2DM affects the transcription machinery regulating monocyte differentiation from GMP versus MDP is presently unknown and warrants further investigation.

The continuum of monocyte transcriptome expression patterns under pathological conditions in particular further underscores the heterogeneity of monocytes and suggests that the Ly6C phenotype by itself does not correlate with the activation status of the monocytes, nor does it reflect the complexity of the functional state ^48,51^ ^47^. Using scRNAseq we have identified a total of 8 clusters of monocytes based on unique cluster-defining gene profiles, beyond the conventional monocyte classification scheme. There was an expansion of cell number especially in Ly6C+ inflammatory monocytes in response to pathological conditions.

Two potential mechanisms by which T2DM or stroke may contribute to the diversity of the monocyte subsets and the heterogeneity of monocyte transcriptome. The first mechanism is the change in myelopoiesis in pathological conditions. Previous studies revealed that circulating monocytes are increased in diabetic mice, both in number and proportion, through accelerated myelopoiesis caused by hyperglycemia ^52,53^, which is consistent with the increased proportion of monocytes in the T2DM groups compared to non-diabetic groups as evidenced by scRNAseq and FACS in our study. In the T2DM groups, monocytosis was accompanied by lymphopenia in the peripheral blood, although we have no data on the production, egress or turnover rate of hematopoietic cells in the bone marrow or circulation to pinpoint the precise causes of increased myeloid cells in T2DM. While the function of neutrophil-like monocytes is not entirely understood, the granule proteins expressed by this population ^46^ could potentially increase brain injury after stroke by directly releasing deleterious substances or other inflammatory mediators ^54^. Therefore, monocytosis with expanded neutrophil-like monocytes may contribute to the aggravated brain damage in T2DM. The second mechanism is the change of monocyte subsets and transcriptome under conditions of T2DM or stroke, indicating that monocytes shaped by the pathological blood microenvironment shared transcriptome signature with monocyte-derived macrophages. This high plasticity of monocytes was seen not only in Ly6C+ subsets but also in Ly6C-subsets in this study. Ly6C-monocytes are less proliferative and remain in the circulation longer than Ly6C+ monocytes ^55,56^. Ly6C-cells reportedly arise from Ly6C+ monocytes in a Nur77-dependent process involving transcription factors Cebpb and Klf2 ^56,57^.

### Diabetic and stroke effect on monocyte transcriptome

#### Suppression of anti-inflammatory genes and activation of pro-inflammatory genes

Monocytes are a heterogeneous population with anti-inflammatory or pro-inflammatory features depending on their stage of differentiation and on the mechanism by which they are activated ^58^. Monocytes in the T2DM or stroke groups are devoid of the steady-state monocyte subset expressing anti-inflammatory genes such as *Cebpb, Klf2*, and *Jund*. Instead, T2DM upregulated pro-inflammatory genes such as *Chil3, S100a8, Lcn2, Slpi* and *Wfdc21*, and stroke further increased the expression of *Hbb-bs* in db/db mice. *Hbb-bs*, encoding hemoglobin beta subunit adult s chain, is a neuroprotective gene reportedly induced in neurons ^59^ but downregulated in astrocytes after MCAO in the brain ^60^. The alpha globin (Hba) is known to restrict nitric oxide release from vascular endothelial cells, although the copy number of which was not associated with increased stroke risk ^61^. Intracerebral hemorrhage was found to increase Hba and Hbb expression in the neurons to buffer the heme released during clot resolution ^62^. The expression or function of heme-making genes *Hba, Hbb, Fech, Fam 220a* and *Alas2* genes in monocyte subsets is unclear, especially when *Hbb-bs* expression was increased after stroke in db/db monocytes. It is possible that stroke induces the release of immature erythroblasts into circulation, although some transcriptome studies regarded them as RBC contamination ^63,64^.

Suppression of pro-survival genes: Cebpb has a context-specific role in neuroprotective polarization of macrophages ^44,65^. Klf2 is an essential regulator of endothelial functions and its suppression is associated with high glucose-induced endothelial dysfunction leading to atherogenesis ^66^. Also, Klf2 is known as an anti-inflammatory factor and negative regulator of monocyte activation that inhibits NFκB, and is suppressed in chronic inflammation ^67,68^. Jund is known to blunt ischemia/reperfusion-induced brain injury via suppression of IL-1β ^69^. Although Cebpb and Klf2 are key transcription factors involved in the differentiation of Ly6C+ to Ly6C-cells from BM to peripheral blood, reduced gene expression of which did not result in reduced Ly6C-population in the db/db mice per scRNAseq and FACS data. Ybx1 and Elf3 are pro-survival transcription factors known to stabilize *bcl2* mRNA ^70,71^ and regulate cell cycle and growth ^72^, respectively. *FAM49B* also affects cell proliferation^73,74^.

S100a8/9 drives the production of neutrophils and monocytes during hyperglycemia and also promotes monocyte and neutrophil recruitment into the arterial wall, potentiating the development of stroke through atherogenesis ^52,75,76^. The upregulation of these genes in the T2DM monocytes is consistent with their biological function.

#### Suppression of IFN-related genes in diabetes

Pathway analysis revealed that the down-regulated pathways in T2DM groups were mostly relevant to inflammation, in which the suppression of Jak-Stat signaling, and MHC antigen presentation can directly lead to the state of immunosuppression and infection-prone. The suppressed IFNγ-inducible genes in the T2DM groups such as *Stat1* and *Gbp2* are central mediators of IFNγ-induced gene expression and critical players in the immune response against viruses, bacteria and parasites ^77,78^. Consistent with this notion, mice deficient in Stat1 were shown to die within eight weeks of birth from viral infections ^79^. The fact that IFN-related genes were down-regulated specifically in T2DM groups implicates a role for deficient anti-infection capacity in the etiology of diabetes ^80,81^.

Downregulated IFN signaling could also impact TLR signaling pathway, which has been shown to play a key role in neuroprotection related to preconditioning effects ^16,18,82^. In particular, Irf3 and Irf7 were reported to be necessary for the preconditioning effects through TLR signaling ^18^. Our transcriptome data revealed that a suite of interferon stimulating genes (ISGs) were strongly suppressed in the T2DM mice after stroke compared to control mice, suggesting that impaired neuroprotection can be one of the detrimental effects of T2DM on stroke outcome. Western blot analysis confirmed that key players in the IFN pathways were also downregulated in the brain in the T2DM mice after stroke and after LPS PC, which is consistent with the failed neuroprotection in diabetic mice following LPS-induced preconditioning. Our data further suggest that similar to aging, T2DM may dampen the host tolerance to ischemic injury.

### Detrimental effect of diabetes on stroke outcome

The current transcriptome data provide mechanistic insights into the detrimental effects of T2DM on stroke outcome. First, immunosuppression can worsen functional outcome in T2DM with stroke. Acute ischemic stroke is known to induce immunosuppression and increased susceptibility to infections ^83,84^, while infection is an independent predictor of neurological deterioration in stroke ^85^. Furthermore, infection after stroke is associated with occurrence of other medical complications, such as gastrointestinal bleeding, decubitus ulcer, deep vein thrombosis, recurrent stroke, and atrial fibrillation ^86^. Reduction in the expression of genes in the IFN pathways and antigen presentation in db/db mice may underlie the increased risk and significant morbidity and mortality from infections and sepsis among stroke patients with diabetes ^80,87^. These data suggest that stroke can exacerbate T2DM complications in infection, resulting in the poor outcomes for these patients. Therefore, the systemic diabetic effects of immunosuppression leading to an infection-prone state should not be overlooked.

The second potential lethal effect is aggravated brain damage caused by increased malfunctioned monocytes in T2DM. Genes involved in neutrophil degranulation, leukocyte migration and chemotaxis are upregulated in T2DM at a more pronounced level after stroke, may potentiate a more severe neuroinflammation after ischemic stroke. In addition, the balance of pro-inflammatory and anti-inflammatory monocytes/macrophages subpopulations is considered to determine the development of brain damage and functional outcome in stroke ^88–90^, as evidenced by the aberrant polarization of brain macrophages and aggravated stroke outcome in hyperglycemia ^15^. Our data suggest a potential relationship between malfunctioned monocytes with impaired neuroprotective function and aggravated brain damage in T2DM. In this regard, suppressed molecular networks related to IFN and TLR caused by an imbalance of pro-inflammatory and anti-inflammatory gene expression in peripheral monocytes may play a key role in the aggravation of stroke outcomes in T2DM.

The third major devastating effect of T2DM on stroke outcome can be attributed to the overactivation of coagulation and complement related genes found in monocytes. The complement and coagulation systems not only share common ancestral genes, but also interact at the protein levels among immune cells, platelets and endothelial cells ^91,92^ ^93^. For example, thrombin, although not a component of C5 convertase, can cleave C5 and produce C5a-like molecules to promote transmigration of myeloid cells into brain after stroke ^94,95^. A number of complement proteins also promote platelet activation and aggregation ^96–98^, enhancing leukocyte-platelet rolling on endothelium after stroke ^99,100^, which can result in poor cerebral blood perfusion ^26,101,102^, increased netosis and thus increased ischemic injury.

### The impact of IFN signaling defect in immunity

IFN signaling defect in T2DM can have a far-reaching impact in health beyond stroke-caused brain injury and complications. First, it may have ramifications in host response to infection in general, which can be influenced by anti-microbial potential and the capacity in antigen presentation of the immune system. Patients with T2D are well known to be more prone to infection ^103^. The molecular mechanism in fighting infection apparently involves IFN signaling, as supported by a recent genome-wide association studies showing that low expression of *IFNAR2 is* associated with life-threatening COVID-19 ^104^. Corroborating in vitro studies also showed that high glucose suppressed type 1 IFN production in PBMC stimulated by poly I:C ^105^, and that bacteria–infected PBMCs from diabetic patients had decreased IFNγ induction and poor bacterial killing ^106^.

Apart from innate anti-microbial capacity, healthy immunity also relies on effective antigen presentation. Evidence suggests that the constitutive expression of Class II major histocompatibility (MHCII) molecules is restricted to professional antigen presenting cells like macrophages, dendritic cells, and B cells, but can be induced by cytokines ^107,108^. Both constitutive and inducible expression of MHC II genes and most MHC II− related chaperone genes are exquisitely controlled at the transcriptional level by a master regulator called Class II Transactivator (CIITA), while CIITA itself is regulated at both the transcriptional and post-translational levels ^107,109^. Interestingly one major inducer of MHCII gene expression is IFNγ, and the induction of CIITA by IFNγ in macrophages involves STAT1 activation by JAK, JNK and IRF1 in the macrophages ^110^. Thus, the defective IFN signaling in T2DM groups observed by scRNAseq and bulk RNAseq in our study may underlie the reduced expression of MHC II genes and contribute to the diminished antigen presentation capacity, resulting in aberrant immunosuppression.

As an integral part of the processing and trafficking of class II MHC complex, CD74 renders MHCII competent for antigen presentation ^111^. Stimulation of IFNγ has been reported to upregulate the expression of CD74, and MIF-CD74 axis in turn also augments the activation of IFNγ-JAK-STAT signaling ^112–114^. T2DM-associated reduction in CD74 expression correlates with reduced H2 gene expression, which is likely attributed to the IFN signaling defect. However, the complex relationship among the IFNγ level, IFN signaling, reduced expression of CD74 and antigen presentation in T2DM warrants further investigation.

Another adverse consequence of IFN signaling defect following infection is the overproduction of proinflammatory cytokines. Recent studies suggest that the causes of the increased susceptibility of patients with T2DM to severe SARS-CoV-2 cytokine storm is in part attributed to infection-induced loss of SETDB2, a transcription factor positively regulated by IFNβ via the JaK1/STAT3 pathway and an enzyme methylates histone 3 lysine 9, leading to increased production of inflammatory cytokines (IL-1β, TNFα, and IL-6) in macrophages via increased activity of H3K9me3 at NFkB binding sites on inflammatory gene promoters following infection ^115^. Corroborating results showed that both coronavirus-infected T2D human plasma and diabetic murine macrophages have reduced levels or expression of *IFNB1* than their non-T2D infected controls, while exogenous administration of IFNβ can reverse inflammation, particularly in diabetic macrophages via SETDBs ^115^.

Although our transcriptome data have revealed novel information in the diversity of peripheral monocytes and mapped distinct biological pathways specific to T2DM, a major limitation of our study is that RNAseq cannot provide information in transcriptional regulation of gene expression such as alternative splicing of genes or epigenetic regulation of gene transcription. Established evidence suggests that the latter mechanism is responsible for regulating IFN/Stat1 signaling. Acetylation of STAT1 inhibits IFN-dependent STAT1 phosphorylation and nuclear translocation, while altering the histone acetyltransferases (HATs) and histone deacetylases (HDACs) activity ratio changes STAT1 activity ^116^. Unlike insulin resistance in type 2 diabetes, viral infection is implicated in the development of islet autoimmunity and type 1 diabetes in which IFN response plays a key role ^117–121^. A study found that macrophages from T1D mice exhibited increased activity of histone acetyltransferases (HAT) and decreased histone deacetylases (HDAC) activity, suggesting that a strict balance between HDAC and HAT activities maintains homeostatic STAT1 expression ^122^. Nonetheless, the epigenetic and metabolic reprogramming of the innate immune cells by chronic T2DM, a phenomenon akin to “trained immunity” ^123,124^, may underlie the monocyte transcriptome signature of hyperinflammatory states seen in the current study. Whether and how the imbalance of HDAC and HAT activities is involved in the downregulation of IFN/STAT1 pathways in the current T2DM model need further investigation.

Second, by focusing on monocyte transcriptome our study does not provide a full view of how T2DM or stroke alters peripheral immunity. Although scRNAseq and FACS data suggest that T2DM also has a great impact on lymphocyte and granulocyte populations, the current study focuses on monocyte transcriptome due to the constraint of cell isolation protocol. Our ongoing studies suggest that T2DM increases neutrophil transmigration into the brain after ischemic stroke, future study in how T2DM alters granulocytes migration and transcriptome is crucial in understanding the detrimental effect of T2DM on inducing netosis and exacerbating ischemic brain injury. Third, our RNAseq experiments were conducted at one time point after stroke which restricts knowledge in the evolution of monocyte gene expression, although it does not impact our understanding of the effect of T2DM on immunity.

In summary, we demonstrated the transcriptome changes in peripheral monocytes under the conditions of T2DM and ischemic stroke using scRNAseq and bulk RNAseq, contributing to their diversity. Expansion in activated pro-inflammatory monocytes and drastic reduction in anti-inflammatory monocytes at steady state can lead to immunosuppression and aggravated brain damage in T2DM with stroke through suppressed IFN-related pathways. The imbalance in the innate immune system caused by T2DM and stroke deregulates the inflammatory process and exacerbates secondary injury, leading to the poor outcome of stroke patients with T2DM. These findings provide novel insights into the underlying mechanism of the adverse effect of T2DM on native immunity and response to stroke.

## Author Contributions

SO performed MCAO and RNA extraction for bulk RNAseq, FACS, data analysis and contributed to the writing of the manuscript. YK carried out pathway analysis of bulk RNAseq, meningeal gene expression and analysis, and MCAO for western blot analysis. YS performed stroke outcome studies for LPS PC. JF performed neurological evaluations and western blot analysis. HZ isolated brain extracts and performed western blot analysis. AK performed cell isolation and MCAO for scRNAseq. WLE performed bioinformatic analysis for bulk RNAseq. HE provided interpretation and clinical implication. CLH provided interpretation and assisted manuscript editing. DA conducted bioinformatics for scRNAseq and enrichment analysis and offered critical interpretation of the study. JL conceived the study design, planning and supervision of the experiments, interpretation of results and writing of the manuscript. All authors have read and agreed to the published version of the manuscript.

## Acknowledgement

This work was supported by NIH grant R01NS102886 (JL), VA Merit Award 2I01BX003335 (JL), Research Career Scientist award 2IK6BX004600 (JL), VA Merit Award I01BX002690 (CLH), AHA 24POST1187683 (HZ).

## Institutional Review Board Statement

The study was conducted in accordance with the Guide for Care and Use of Laboratory Animals issued by the National Institutes of Health and approved by San Francisco Veterans Affairs Medical Center Institutional Animal Care and Use Committee (protocol number 20-012 and 23-013, approved 8-10-2020, 9-6-2023).

## Supplemental Figure/Table Legends

**Figure S1.**
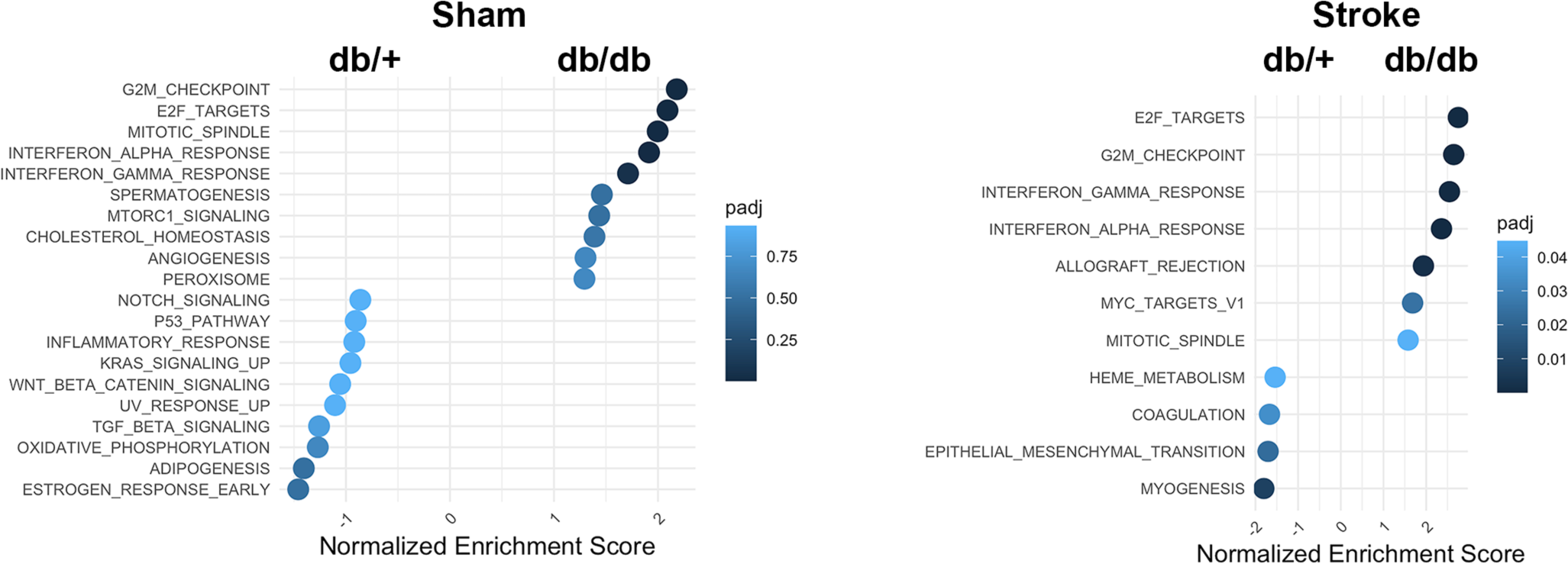
Gene set enrichment analysis based on HALLMARK database. DEGs with FDR less than 0.05 were used as the input genes from peripheral blood monocyte bulk RNA sequencing data and ranked by log2FC. Top 10 pathways were identified and visualized in the dotplot.

**Table S1.**
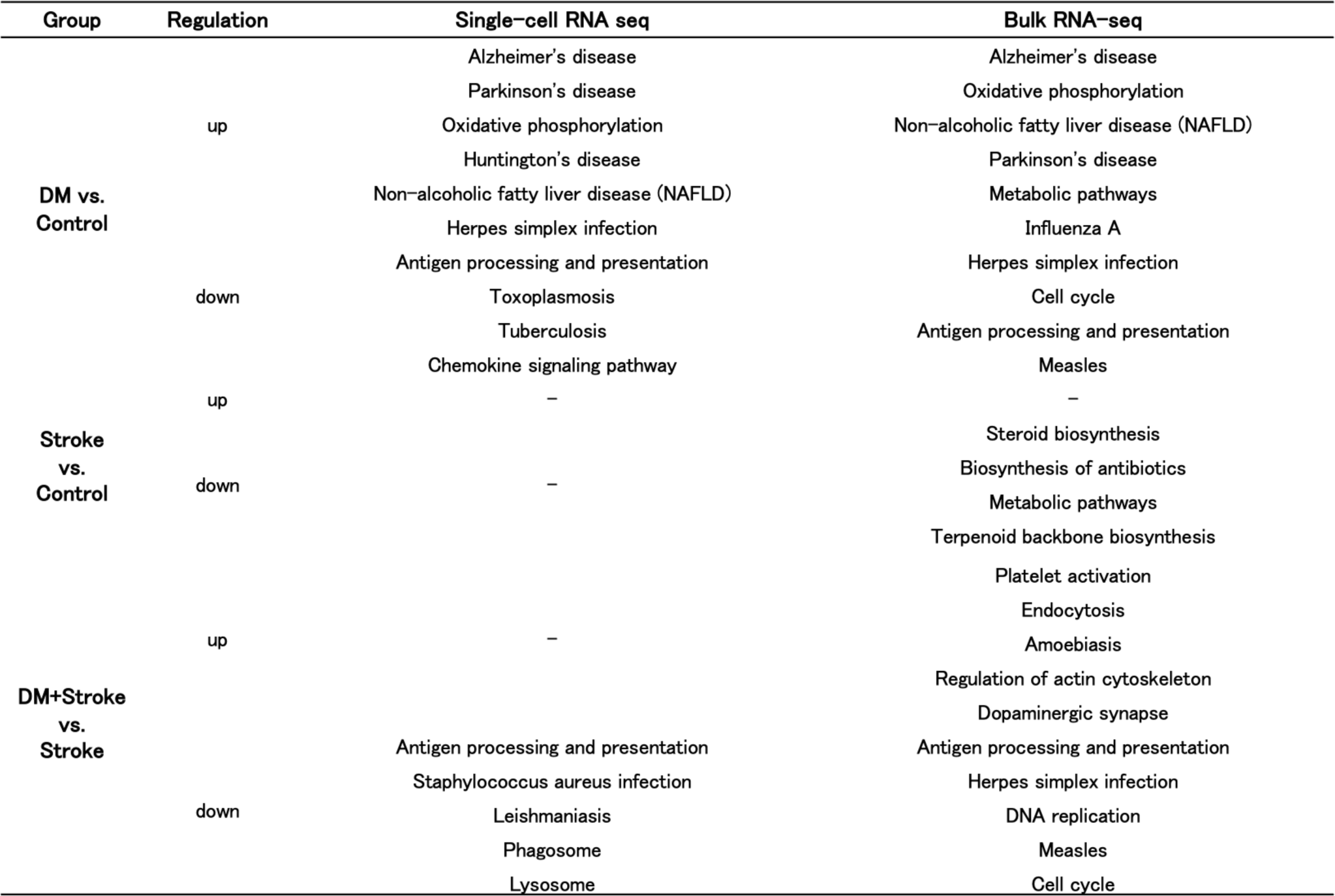
Top five Kyoto Encyclopedia of Genes and Genomes (KEGG) pathways enriched for each group comparison with single cell- and Bulk-RNA sequencing. T2DM groups had down-regulated pathways related to infection such as “herpes simplex infection” and “antigen processing and presentation”. Similarly, bulk RNA-seq analysis showed enriched these pathways in T2DM groups. DEGs related to each pathway are shown in supplementary tables. Specifically, the DEGs in the herpes simplex infection pathway were also involved in sub-pathways of Jak-Stat signaling (such as Stat1), MHC antigen presentation (H2A family), and Toll-like receptor (TLR) signaling (such as Irf7).

| Group | Regulation | Single-cell RNA seq | Bulk RNA-seq |
| --- | --- | --- | --- |
| **DM vs. Control** | up | Alzheimer's disease | Alzheimer's disease |
|  |  | Parkinson's disease | Oxidative phosphorylation |
|  |  | Oxidative phosphorylation | Non-alcoholic fatty liver disease (NAFLD) |
|  |  | Huntington's disease | Parkinson's disease |
|  |  | Non-alcoholic fatty liver disease (NAFLD) | Metabolic pathways |
|  | down | Herpes simplex infection | Influenza A |
|  |  | Antigen processing and presentation | Herpes simplex infection |
|  |  | Toxoplasmosis | Cell cycle |
|  |  | Tuberculosis | Antigen processing and presentation |
|  |  | Chemokine signaling pathway | Measles |
| **Stroke vs. Control** | up | – | – |
|  | down | – | Steroid biosynthesis |
|  |  | – | Biosynthesis of antibiotics |
|  |  | – | Metabolic pathways |
|  |  | – | Terpenoid backbone biosynthesis |
| **DM+Stroke vs. Stroke** | up | – | Platelet activation |
|  |  | – | Endocytosis |
|  |  | – | Amoebiasis |
|  |  | – | Regulation of actin cytoskeleton |
|  |  | – | Dopaminergic synapse |
|  | down | Antigen processing and presentation | Antigen processing and presentation |
|  |  | Staphylococcus aureus infection | Herpes simplex infection |
|  |  | Leishmaniasis | DNA replication |
|  |  | Phagosome | Measles |
|  |  | Lysosome | Cell cycle |

**Table S2.**
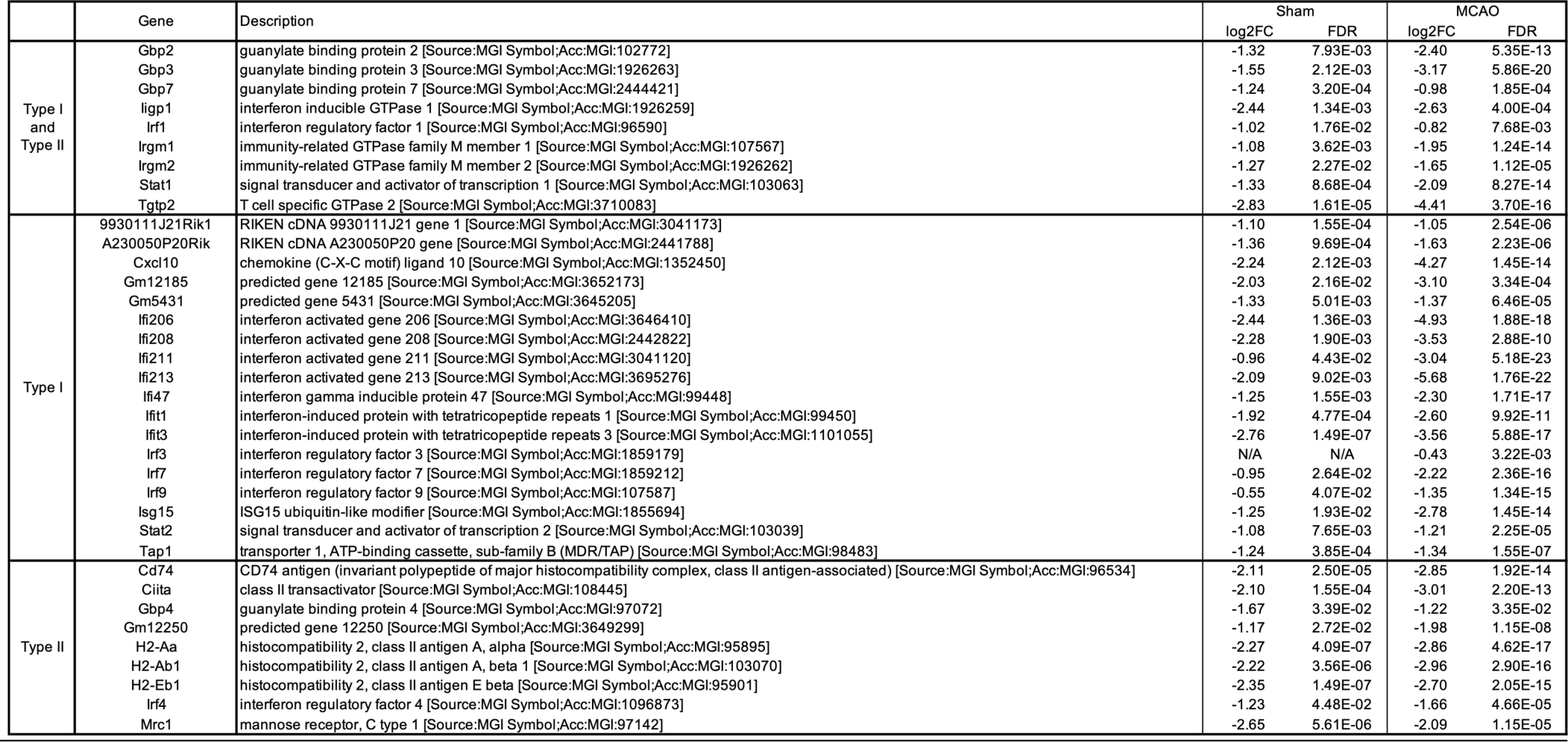
Representative downregulated DEGs in the type 1 and type 2 IFN pathways comparing db/db to *db/+* monocytes in sham and MCAO groups. Fold change in Log 2 scale and adjust P values are shown for each gene.

**Table 3.**
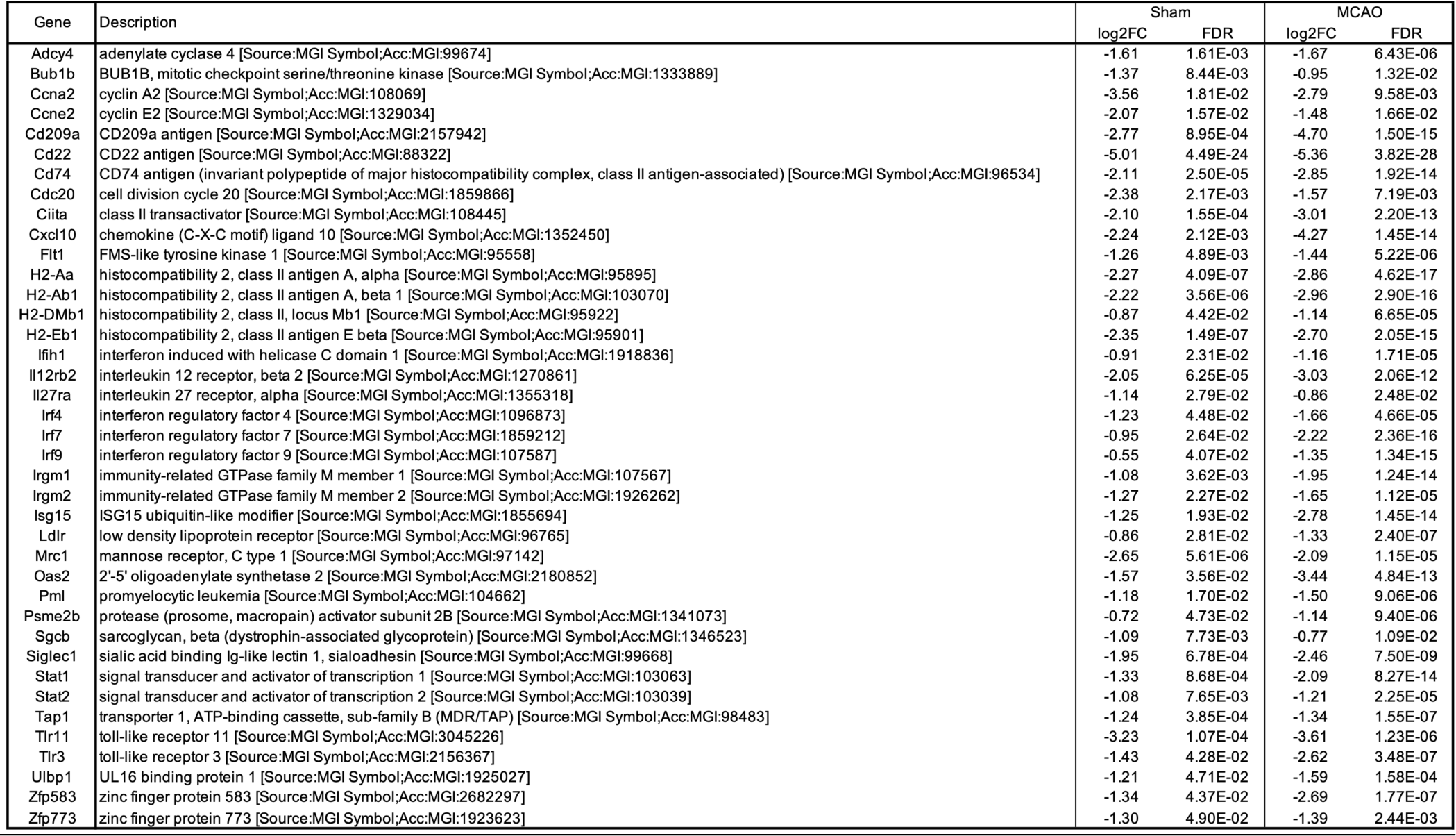
Representative downregulated DEGs relevant to antigen presentation and class II MHC comparing *db/db* to db/+ monocytes in sham and MCAO groups. Fold change in Log 2 scale and adjust P values are shown for each gene.

**Table 4.**
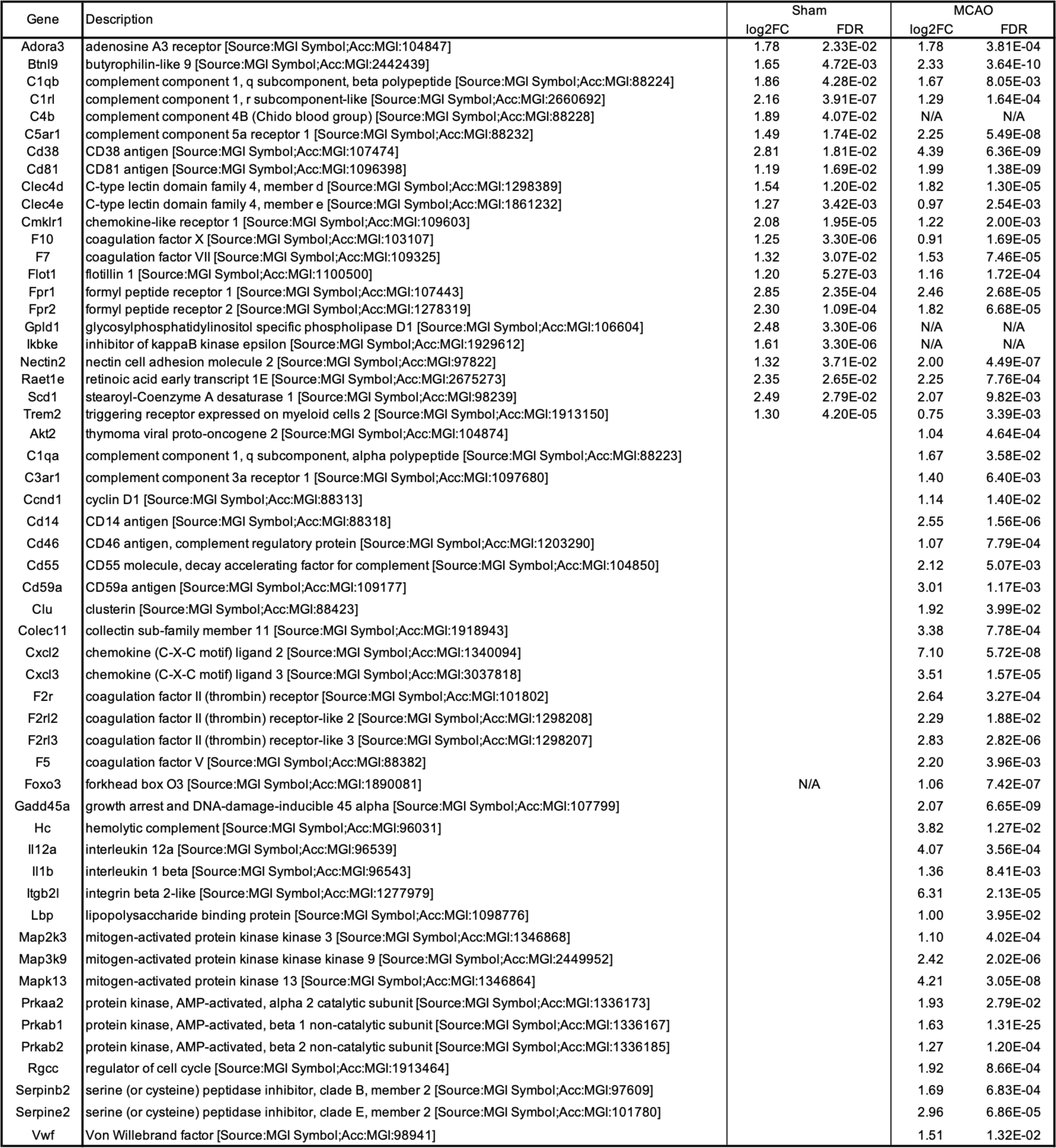
Representative upregulated DEGs in the complement and coagulation cascades comparing db/db to db/+ monocytes in sham and MCAO groups. Fold change in Log 2 scale and adjust P values are shown for each gene.

